# The mechanism of ribosomal recruitment during translation initiation on Type 2 IRESs

**DOI:** 10.1101/2025.06.11.659010

**Authors:** Sayan Bhattacharjee, Irina S. Abaeva, Zuben P. Brown, Yani Arhab, Hengameh Fallah, Christopher U. T. Hellen, Joachim Frank, Tatyana V. Pestova

## Abstract

The encephalomyocarditis virus (EMCV) IRES and other Type 2 IRESs comprise domains H-L and specifically interact with eIF4G/eIF4A through their essential JK domain. However, the JK domain is not sufficient for IRES function, which also requires the preceding domain I of unknown function. To identify interactions that drive ribosomal recruitment of eIF4G/eIF4A-bound Type 2 IRESs, we determined the cryo-EM structure of 48S initiation complexes formed on the EMCV IRES. It revealed that the apical domain I cloverleaf contacts ribosomal proteins uS13 and uS19 via its Id subdomain and that the essential GNRA tetraloop in subdomain Ic interacts directly with the TψC domain of initiator tRNA. Functional assays supported the exceptional role of these interactions for initiation on this IRES. The strong conservation of primary and secondary structures of the apex of domain I among Type 2 IRESs suggests that the reported interactions are a common essential feature of them all.

## INTRODUCTION

Initiation on most cellular mRNAs occurs by the 5’end-dependent ribosomal scanning mechanism and involves multiple eukaryotic initiation factors (eIFs)^1^. A 40S ribosomal subunit binds an eIF2•GTP/Met-tRNA_i_^Met^ ternary complex, eIFs 3, 1 and 1A to form a 43S preinitiation complex (PIC). Attachment of 43S PICs to the capped 5’-end of mRNA is mediated by eIF4F (comprising the cap-binding protein eIF4E, the RNA helicase eIF4A, and the scaffold protein eIF4G which also binds to eIF3), eIF4A and eIF4B. eIFs 4A/4B/4F unwind the cap-proximal region of mRNA allowing attachment of 43S PICs that is also facilitated by the eIF4G-eIF3 interaction. 43S PICs subsequently scan to the first AUG in a favorable context, where they form 48S initiation complexes (ICs) with codon-anticodon base-pairing in the ribosomal P site. eIF1, in cooperation with eIF1A, monitors the fidelity of initiation. Start codon recognition leads to eviction of eIF1, eIF5-induced hydrolysis of eIF2-bound GTP and Pi release, committing the 40S subunit to the initiation codon. Subsequent joining of a 60S ribosomal subunit and release of eIFs are mediated by eIF5B, which reorients the acceptor end of Met-tRNA_i_^Met^ to allow subunit joining^2,3^ and then hydrolyzes GTP and dissociates from assembled 80S ribosomes.

Viruses have evolved diverse mechanisms to sustain translation of their mRNAs in infected cells after innate immune responses have been activated. One approach employs an internal ribosomal entry site (IRES), which is a *cis*-acting RNA element that promotes end-independent ribosomal recruitment to an internal location in mRNA^4^. There are four principal classes of IRES: Type 1 (e.g. poliovirus), Type 2 (e.g. encephalomyocarditis virus (EMCV)), Type 4 (e.g. hepatitis C virus (HCV)) and Type 6 (e.g. cricket paralysis virus (CrPV) and Halastavi árva virus (HalV)) (nomenclature according to Ref.5). These classes employ distinct mechanisms for ribosome recruitment that are nevertheless all based on specific non-canonical interactions with canonical components of the translation apparatus and require only a subset of canonical eIFs. Thus, Type 4 IRESs bypass the requirement for eIFs 4A, 4B, 4F and eIF3 by binding directly to the platform of the 40S subunit, usurping the eIF3 binding site^2,6,7^, whereas type 6 IRESs bind to either the ribosomal A site (CrPV^8,9^) or to the P site (HalV^10^) and do not require any eIFs or initiator tRNA. On Type 4 IRESs, Met-tRNA_i_^Met^ can be recruited not only by eIF2 but also by eIF5B or even eIF2D^11–13^. Importantly, cryo-EM studies have been indispensable in gaining molecular insights into the mechanisms of initiation on Type 4 and Type 6 IRESs^2,7,9,10,14–21^.

Type 2 IRESs were initially discovered in members of *Picornaviridae*^22,23^ and were recently also identified in *Caliciviridae*^24,25^. They are ∼450nt-long and have five principal domains (H-L), with a Yn-Xm-AUG motif at their 3′-border in which a Yn pyrimidine tract (n = 8-10 nt) is separated by a spacer (m = 18-20 nt) from an AUG triplet that acts as the initiation codon (AUG_834_ in the EMCV IRES; Figure S1). *In vitro* reconstitution has shown that initiation on Type 2 IRESs requires eIF2, eIF3, eIF4A and the central domain of eIF4G, that it is stimulated by eIF4B and the pyrimidine tract-binding protein PTB (or its neuronal paralogue nPTB), an IRES *trans*-acting factor (ITAF), and that eIF1 and eIF1A synergistically enhance the fidelity of initiation codon selection^26–30^.

Each domain in type 2 IRESs has a conserved structure and contains highly conserved sequence motifs, mostly at peripheral locations, that are critical for function. PTB binds to oligopyrimidine sequences in different domains and is thought to stabilize the IRES in an active conformation^31,32^. The only known essential interaction with canonical components of the translation apparatus for Type 2 IRESs is that of their JK domains with eIF4G’s central domain^26,27,33–37^ (Figure S1). eIF4G binds to highly conserved elements of the J-K domain in a manner that is enhanced by eIF4A^36–38^, and together, they induce conformational changes at the IRES’s 3’ border, in the region that enters the mRNA binding cleft of the 40S subunit^39^. The JK domain is essential, but it is not sufficient for IRES activity, which also requires domain I^22,23,40,41^. The function of domain I remains obscure. It consists of a long, irregular stem and an apical double cross structure with conserved loops, including a GNRA tetraloop. GNRA tetraloops adopt a specific U-turn structure which is strongly stabilized by networks of hydrogen bonding and base-stacking within the loop^42–44^. This structure is characterized by a sharp turn of the phosphate backbone between G and the second nucleotide and the last three bases being sequentially stacked on each other. This structure exposes the Watson-Crick base-pairing edge of these three bases, enabling GNRA tetraloops to engage in long-range tertiary interactions^45^. GNRA tetraloops are consequently often involved in interactions with other RNA elements (e.g. in ribosomal RNAs) or proteins, playing a key role in RNA folding and function. In Type 2 IRESs, GNRA loops are also critical for the IRES activity, but their exact function and potential binding partners remain unknown^46–52^.

The facts that the JK domain is not sufficient for EMCV IRES function and that initiation on it does not depend on eIF4G’s eIF3-binding domain^38,53^ raise the question of which interactions are responsible for recruiting an eIF4G/eIF4A-bound Type 2 IRES to a 43S PIC. To resolve this question, which is of paramount importance for this mechanism of initiation, we determined the cryo-EM structure of 48S complexes assembled on the EMCV IRES.

## RESULTS

### The structure of the 48S complex assembled on the EMCV IRES

To identify the interactions that are responsible for the recruitment of Type 2 IRESs to 43S preinitiation complexes, we determined the cryo-EM structure of the 48S initiation complex assembled on the EMCV IRES. 48S complexes were reconstituted *in vitro* on EMCV IRES-containing mRNA (nt 315-862) using individual 40S ribosomal subunits, Met-tRNA_i_^Met^, initiation factors eIF2, eIF3, eIF4A, eIF4B, eIF4F, eIF1, eIF1A, as well as the EMCV-specific ITAF nPTB. *In vitro* reconstitution done using purified components ensures the assembly of complexes of known composition. Accurate assembly of 48S complexes on the IRES was verified by primer extension inhibition, which involves extension by reverse transcriptase of a primer base-paired to the mRNA. mRNA-bound ribosomes arrest primer extension, yielding toeprints at their leading edge that can be located on sequencing gels. 48S complexes yield stops 15, 16 and 17 nt downstream of the first nucleotide (^+^1) of the initiation codon. To increase homogeneity, AUG_826_ (which is weakly utilized during *in vitro* reconstitution^28^) was mutated so that 48S complexes assembled exclusively at the authentic start codon AUG_834_. To avoid potential cryo-EM artifacts, chemical cross-linking was not applied following 48S complex assembly.

Grid preparation, data collection and image processing are described in *Methods*. Cryo-EM grids were imaged at 300 kV producing high-resolution micrographs with easily identifiable 40S ribosomal particles (Table S2). From initial 3D classification (see Methods and Figure S2) four classes of interest were selected: Class I (134,626 particles), Class II (98,748 particles), Class III (89,242 particles), and Class IV (72,526 particles). All four classes exhibited densities corresponding to the 40S ribosomal subunit, eIF1A and the IRES bound to the inter-subunit face of the 40S head. Classes II-IV also contained density for eIF1, whereas not fully resolved Class I contained additional density for eIF2, Met-tRNA_i_^Met^, and blurry density for eIF3. Subsequent focused classification on the Class I population revealed a refined class, designated Class Ia, containing 59,153 particles with well-resolved density for the 40S, eIF2, eIF1A, Met-tRNA_i_^Met^, eIF3 and the IRES. Additionally, Classes II–IV that could correspond to initial binding of the IRES to 40S subunits differed from one another in the extent of the rotation of the 40S subunit head and in repositioning of the IRES (Figure S2B), which might represent conformational changes that would gradually open the P site and reduce potential steric clashes for spatial accommodation of Met-tRNA_i_^Met^, eIF2α and the IRES. The smaller particle population observed in Class Ia compared to the other classes likely reflects dissociation of eIF3 from 48S complexes during grid preparation because 48S complexes did not undergo chemical cross-linking. The lack of cross-linking could also account for the absence of eIF4F/eIF4A in our structure^54^. Subsequent refinement of density maps from Classes Ia–IV yielded structures with an approximate global resolution of 3.2Å according to FSC curves (see Methods; Figures S2-S3).

In the final model (Figures 1A-C), 48S complexes showed density corresponding to canonical components of initiation complexes i.e. the 40S subunit, P-site Met-tRNA_i_^Met^ base-paired with the initiation codon and bound to the eIF2ψ subunit, E-site eIF2α subunit, A-site eIF1A, the five-lobed PCI/MPN core of eIF3 bound to its conventional site at the solvent side of the 40S subunit, as well as an additional density for the IRES (red) bound to the head of the 40S subunit and interacting with ribosomal proteins uS13 and uS19 (blue and green, respectively) as well as with the P-site Met-tRNA_i_^Met^ (magenta), and also extending away from the ribosome head.

**Figure 1.**
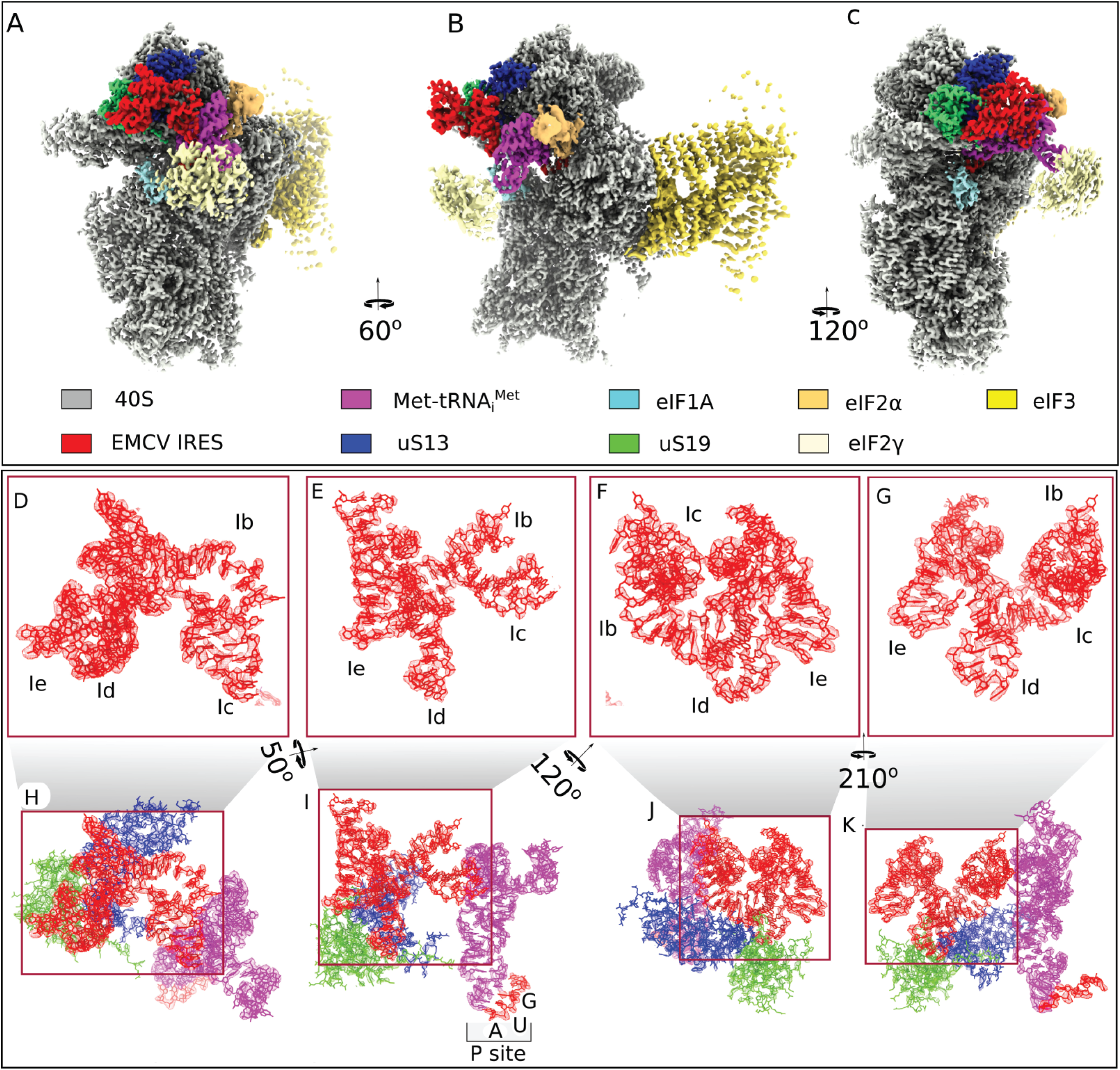
Translation initiation complex on the EMCV IRES. (**A**–**C**) Cryo-EM map of the 48S initiation complex assembled on the EMCV IRES, shown in different orientations. The 40S ribosomal subunit, ribosomal proteins uS13 and uS19, associated initiation factors, Met-tRNA_i_^Met^ and EMCV IRES are shown in distinct colors and are labeled accordingly. (**D**–**G**) Zoomed-in views of the EMCV IRES density map (rendered at 70% surface transparency) superimposed with the atomic model, displayed in different orientations. The individual domains of the IRES are indicated in these views. (**H**–**K**) Zoomed-in views of the cryo-EM density map, with superimposed atomic models, showing the key interactions between the EMCV IRES (red) and initiator tRNA (magenta), uS13 (blue) and uS19 (green). The densities of the different components are color-coded using the ChimeraX surface color tool^67^. Panel **I** also shows interaction between the start codon (AUG) of the mRNA and the anti-codon of the initiator tRNA at the P site of the 40S subunit.

Due to the high resolution of the IRES we could identify contiguous strands of mRNA and classify nucleotides within those strands as either purines (R) or pyrimidines (Y) based on the size of their density. To determine which region of the IRES was bound to the 40S subunit we chose the highest-resolution portion, a loop bound in the pocket formed between alpha helix 2-3 of ribosomal protein uS13 and the beta sheet of uS19 (Figure S4). This region of the IRES consists of 13 nucleotides beginning with a clearly identifiable purine, followed by a pyrimidine, two purines, seven nucleotides in a loop and two pyrimidines. Using the presumptive identity of these nucleotides we searched the IRES for candidate regions beginning with the smallest identifiable sequence (viz. RNRR) and found 39 possible locations. The IRES density clearly showed a 7nt-long loop linking the final two purines of our search term (Rn**RR**) to two facing nucleotides. We therefore eliminated any region that could not accommodate such a loop from further consideration and were left with two candidates, in domain H (^414^AGGG_417_) and in subdomain Id (^562^GCGG_565_). Examination of the IRES density suggests that an additional pyrimidine is present at either end of the search term (**Y**RNRR**Y**) and by examining each separately (e.g., RNRRY and YRNRR) we eliminated domain H as a possibility due to the length of the loop. We exhaustively increased the length of the search term as well as the direction (e.g., 5ʹ to 3ʹ or 3ʹ to 5ʹ) (Table S2) and in all cases the sequence in subdomain Id was identified as the only feasible candidate. Although the density of the loop suggested an unambiguous length of seven nucleotides connecting the two purines (^564^GG_565_) to their base-pairing pyrimidines (^573^CC_574_) we also varied the length during our search to minimize bias and found no possibilities that conformed with the predicted secondary structure of the IRES. Full details of the parameters that we used to identify possible candidate sequences are given in the *Methods*. Once we had determined the identity of the _561_TGCGGCCAAAAGCC_574_ region, we were able to build subdomains Ib, Ic, Id, and Ie that span 83 nucleotides in the apex of domain I (_518_CC..GG_600_) accounting for 17% of the IRES (Figures 1D-G and S1), and that interact with Met-tRNA_i_^Met^, uS13 and uS19 (Figures 1H-K).

To verify the assignment of cryo-EM density to the apical region of domain I and to confirm its identical position in 48S complexes in solution, we performed directed hydroxyl radical cleavage of the IRES from the surface of eIF1A in *in vitro* assembled 48S complexes and also identified the regions of the IRES that are protected from RNase T1 digestion in initiation complexes. We observed hydroxyl radical cleavage at the junction of Id and Ie subdomains (CCA_573-575_) from two positions in the C-terminal tail of eIF1A (D_132_ and D_137_) (Figures S5A-B). The modeled position of these IRES nucleotides is in very close proximity to 18S rRNA nucleotides that are cleaved from the same positions of eIF1A in 43S preinitiation complexes^55^ (Figure S5C), confirming the assignment. Protection from RNaseT1 cleavage of G_549_ in subdomain Ic and of G_585_ in subdomain Ie was observed in reaction mixtures for 48S complex formation (Figure S6A, left panel; Figure S6C) and in assembled, sucrose density gradient purified 48S complexes (Figure S6A, right panel; Figure S6C), which is also consistent with the cryo-EM structure. In addition, we observed protection of G_474_ in the central stem of domain I, an element that was not seen by cryo-EM. Protection of G_775_ and G_761_ in domain JK (Figures S6B-C) corresponded to the previously reported specific binding of eIF4G/eIF4A^27,33^.

### Specific interactions of the EMCV IRES with the 40S subunit and initiator tRNA

Detailed analysis of the model revealed that the apical region of IRES domain I adopts a cloverleaf conformation that allows the Ih2 helix and the A-rich Id loop to interact with ribosomal proteins uS13 and uS19, respectively (Figure 2). Specifically, the α-helix of uS13, composed of charged and polar residues E112, R108, and N105, forms a platform that facilitates docking of the IRES Ih2 helix via its G530 and U561 residues (Figures 2A-F). Further interaction of Q104 and N103 of uS19 with the Id loop helps to support this docking of the IRES to the 40S subunit head (Figures 2G-L).

**Figure 2.**
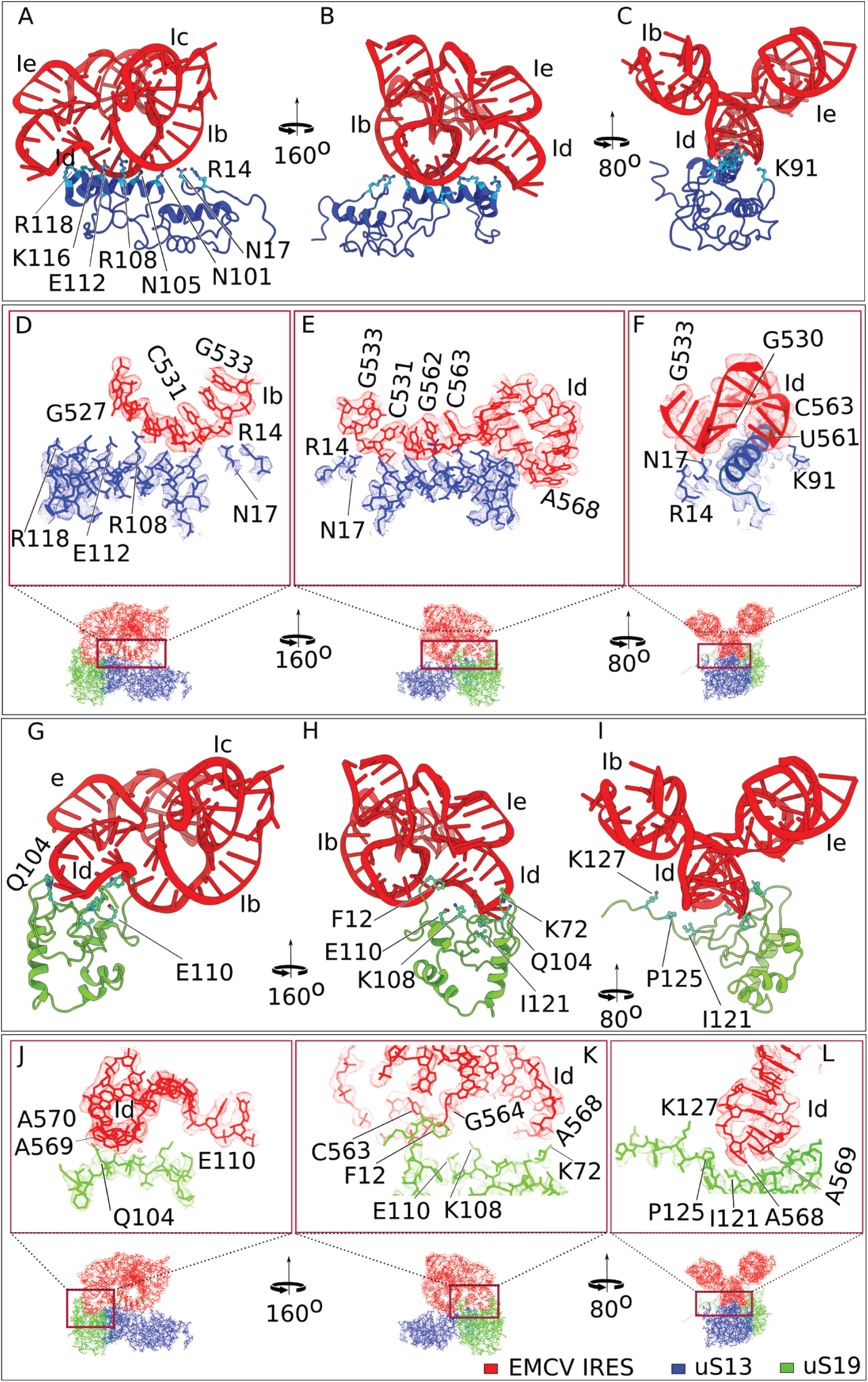
Specific interactions of EMCV IRES domain I with the head of 40S ribosomal subunit. (**A**-**C**) Zoomed-in views of the interaction between protein uS13 (blue) of the 40S subunit and the EMCV IRES, as shown by atomic models in three orientations, with interacting residues labeled. (**D**-**F**) Close-up views of the Id subdomain of the EMCV IRES interacting with uS13, presented in different orientations, with map density rendered at 70% transparency to show the fitting of the atomic model. (**G**-**I**) Interaction between protein uS19 (green) of the 40S subunit and the EMCV IRES (red) shown by atomic models presented in different orientations, with interacting residues labeled. (**J**-**L**) Close-up views of the Ib and Id subdomains of the EMCV IRES interacting with uS19, presented in different orientations, with map density rendered at 70% transparency to show the fitting of the atomic model.

Strikingly, the Ic GNRA loop (GCGA_547-550_), which is essential but whose mechanism of action is unknown, is positioned to interact directly with the initiator tRNA TψC domain (Figures 3A-C). Comparison of the P-site tRNA model in canonical cap-dependent 48S ICs (e.g. Ref. 56) and in the EMCV IRES 48S IC revealed that while binding to the P-site codon the tRNA acceptor domain shifts by 9.7Å from the associated eIF2 to interact with the EMCV IRES (Figure 3B). In the IRES-tRNA interface, bases A550, G549 and C548 in the GNRA loop are connected either via electrostatics or via van der Waals force with G48 and C65 in initiator tRNA (Figures 3D-G). The IRES-Met-tRNA_i_^Met^ interaction is further stabilized by the additional interaction of G546 and A550 in the IRES and U46 in tRNA (Figures 3H-I), where the Met-tRNA_i_^Met^ variable loop containing U46 rotates to interact with IRES.

**Figure 3.**
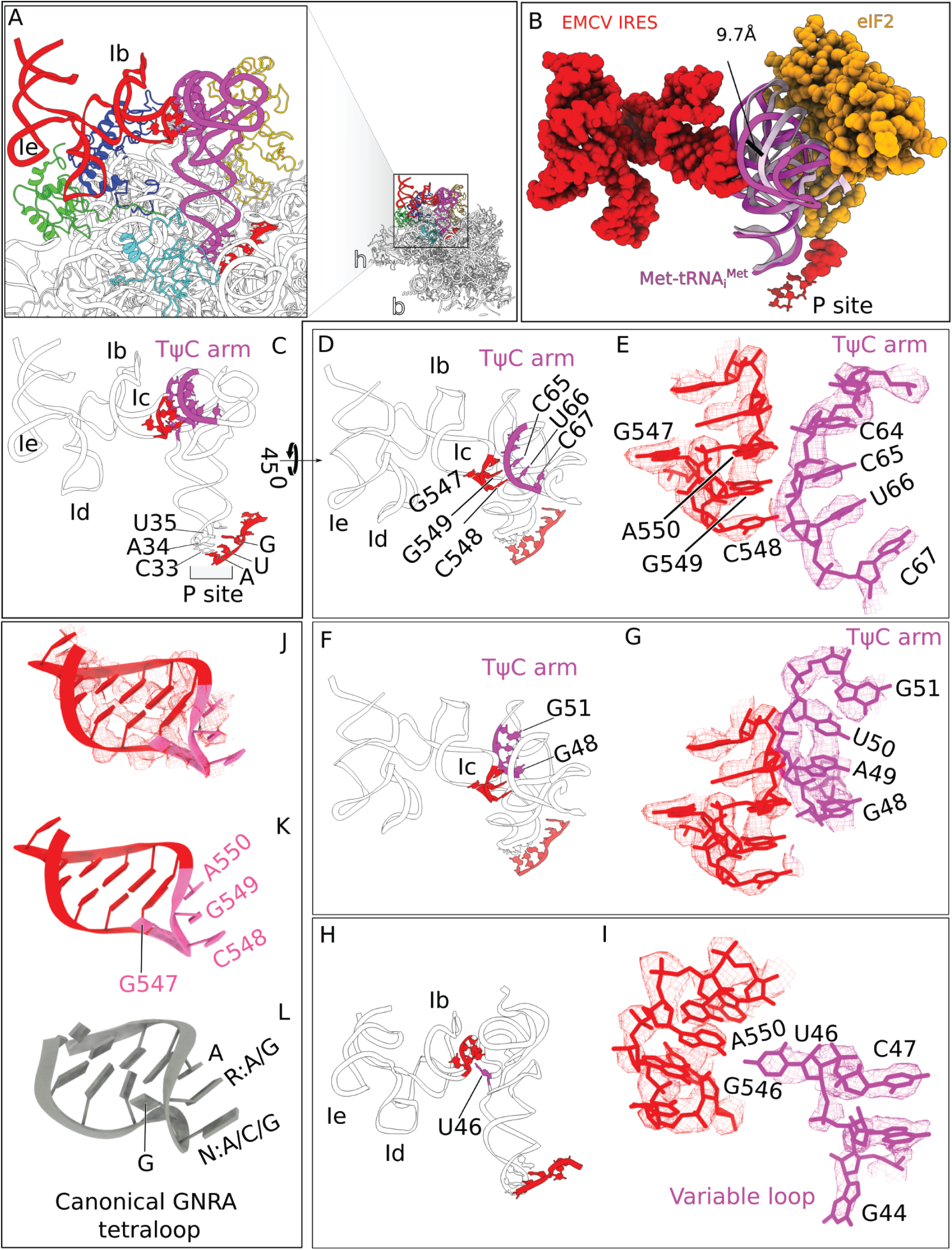
The conformation of the EMCV IRES domain I tetraloop and its interactions with initiator tRNA. (**A**) Close-up view of the atomic model of the 40S subunit head region, showing interactions between the EMCV IRES (red), initiator tRNA (magenta), and 40S subunit proteins uS19 (green) and uS13 (blue). (B) Structural comparison reveals that the acceptor arm of the P-site-bound tRNA undergoes a ∼9.7 Å displacement from its position associated with eIF2 in canonical 48S complexes (PDB id: 8OZ0) (tRNA is in light pink) to a new position where it interacts directly with the EMCV IRES (tRNA is in magenta). (**C**) Interaction between the tetraloop of Ic domain of the EMCV IRES and initiator tRNA, and codon-anticodon recognition between initiator tRNA at the P site and the EMCV initiation codon AUG_834_. (**D-I**) structural details of the EMCV IRES tetraloop regions interacting with initiator tRNA. (**D** and **E**) Interaction is shown by atomic models and by atomic models fitted into Coulomb density (rendered at 70% transparency, and highlighting the front TψC arm), respectively. (**F** and **G**) Same interaction is shown but with a focus on the rear TψC arm. (**H** and **I**) Interaction between the variable loop of initiator tRNA and the tetraloop of the EMCV IRES visualized by atomic models and atomic models fitted into Coulomb density (rendered at 70% transparency). (**J**) conformation of the EMCV IRES tetraloop, shown by the atomic model fitted into Coulomb density (rendered at 70% transparency). (**K** and **L**) Comparison of the tetraloop structure found in the EMCV IRES with canonical GNRA tetraloops (where N can be A/G/C and R is A/G) (PDB: 1ZIF).

Comparison with available structures of GNRA tetraloops revealed that the GNRA tetraloop of the IRES has a similar structure (Figures 3J-L). GNRA tetraloops constitute the largest class of tetraloops and commonly function as determinants of RNA tertiary structure. GNRA tetraloops bind to three types of RNA receptor: the 11nt receptor (which is specific for GAAA loops, which stack on an adenosine platform in the receptor), the IC3 receptor, another asymmetric internal loop present in the IC3 subgroup of self-splicing introns, and the minor groove receptors, which are formed by two helical base-pairs and interact via A-minor type hydrogen bonding with the third and fourth nucleotides of the tetraloop^45,57^. Several variants of the latter type of tetraloop-receptor have been identified, which deviate with respect to the location of the hydrogen bonding interface, in some instances due to rotation of the tetraloop bases away from the receptor. The direct contact between the EMCV mRNA tetraloop and the TψC arm of initiator tRNA observed here represents a structurally distinct and novel pattern of interactions, which contribute directly to complex stabilization and ribosomal positioning of the IRES - features not previously reported in the context of translation initiation.

Mutational analysis of domain I (Figure 4A) confirmed that the structure of its apical region is critical for the IRES function. Thus, disruption of Ih2 and Ih4 abrogated 48S complex formation on the IRES in the *in vitro* reconstituted system, but activity was restored by compensatory mutations (Figure 4B lanes 5-8). Disruption of the Ih1 apical base-pair did not abrogate but strongly reduced the activity of the IRES, whereas the compensatory mutation again restored IRES function (Figure 4B, lanes 3-4). Mutations in Id loop affected IRES activity to different extents, with deletion of two A resides having the biggest effect (Figure 4B lanes 9-12). In accordance with previous reports regarding the importance of the tetraloop for type 2 IRES function^46–48,51,52^, mutations in the GNRA loop also impaired formation of 48S complexes on the EMCV IRES (Figure 4C). Consistent with the intimate interaction of the EMCV IRES with the initiator tRNA, 48S complex formation was also sensitive to post-transcriptional modification of initiator tRNA: *in vitro* transcribed tRNA was less efficient and also promoted a higher level of initiation at the aberrant upstream AUG_826_ independently of the presence/absence of eIF1 and eIF1A (Figure S7A).

**Figure 4.**
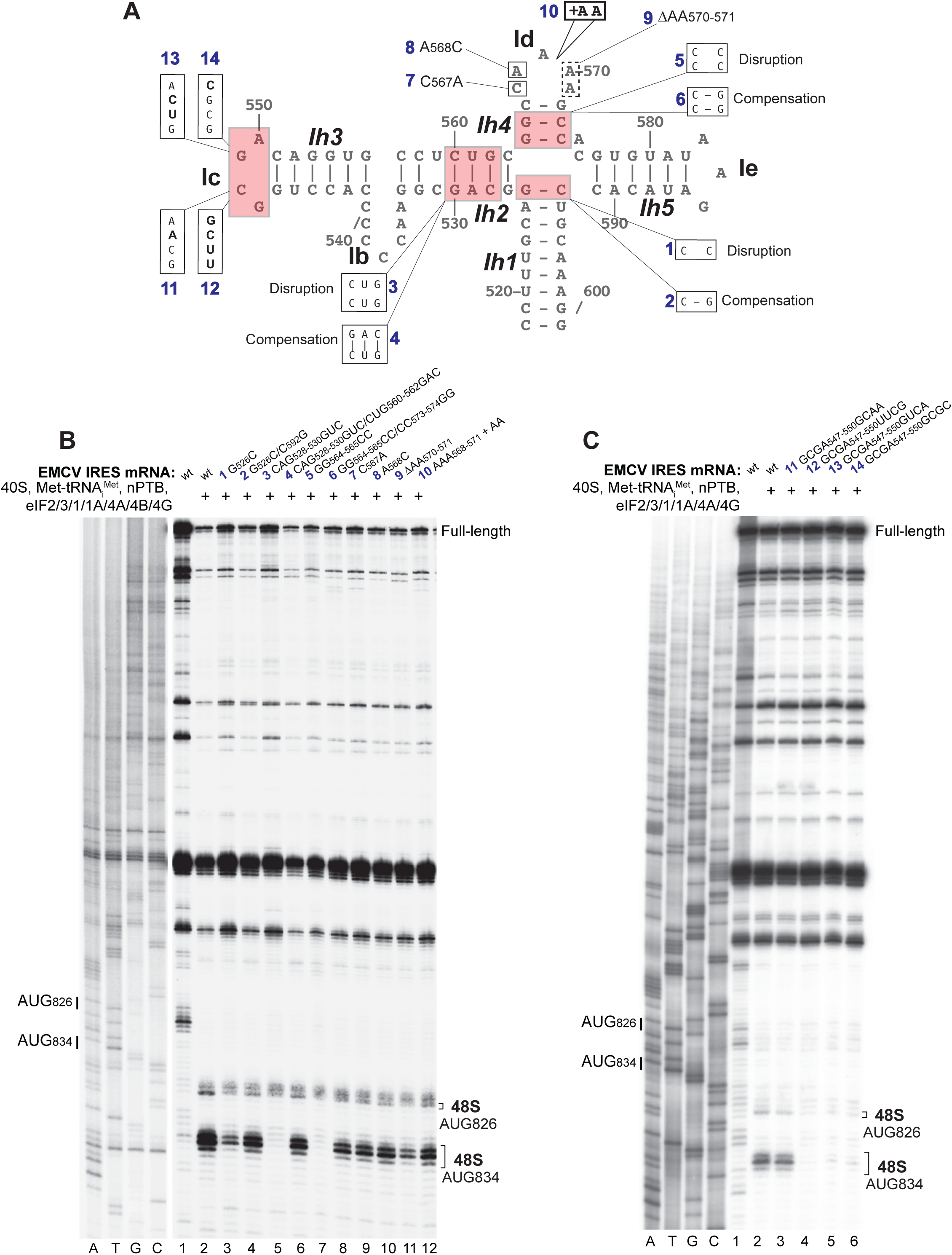
Influence of mutations in the apex of domain I on 48S complex formation on the EMCV IRES. (**A**) Secondary structure model of the apex of domain I showing introduced mutations. (**B-C**) Toe-printing anaysis of 48S complex formation on wt and mutant EMCV IRES mRNAs in the presence of individual 40S subunits, Met-tRNA_i_^Met^, nPTB and indicated eIFs. Lanes C, T, A, and G depict the wt EMCV sequence generated using the same primer. The positions of initiation codons are indicated on the left and of assembled 48S complexes on the right. The division between sequence lanes and lanes 1–12 in (**A**) indicates that these two sets of lanes were derived from the same gel, exposed for different lengths of time.

Interestingly, the interaction of the IRES and uS19 mimics canonical contacts between 28S rRNA of the 60S subunit and uS19 in the classical-1 PRE state^58,59^, suggesting that the IRES evolved to exploit a pre-existing rRNA/rprotein contact. It also implies that the IRES would prevent formation of 80S ribosomes by a steric clash with the 60S subunit (Figures 5A-D) and that displacing of the IRES is a prerequisite for ribosomal subunit joining. Comparison of the position of Met-tRNA_i_^Met^ in 48S complexes assembled on the EMCV IRES with eIF2 with the position of Met-tRNA_i_^Met^ in initiation complexes containing eIF5B^2,3^ indicates a 16.7 Å tRNA shift in the latter (Figures 5E-J). Such a shift, in turn, would result in a steric clash between the tRNA T-loop and C548 of the EMCV GNRA tetraloop (Figure 5K), disrupting the stable tetraloop-tRNA contact and providing a mechanism for IRES displacement required for subsequent subunit joining. Consistently, we found that unlike in the case of HCV-like IRESs, eIF5B could not substitute for eIF2 in 48S complex formation on the EMCV IRES, yielding only trace amounts of initiation complexes (Figure S7B, lanes 3 and 8). eIF2D also could not replace eIF2 (Figure S7B, lanes 4 and 9) suggesting that like eIF5B, it also could not ensure the position of tRNA compatible with its interaction with the GNRA loop of the IRES.

**Figure 5.**
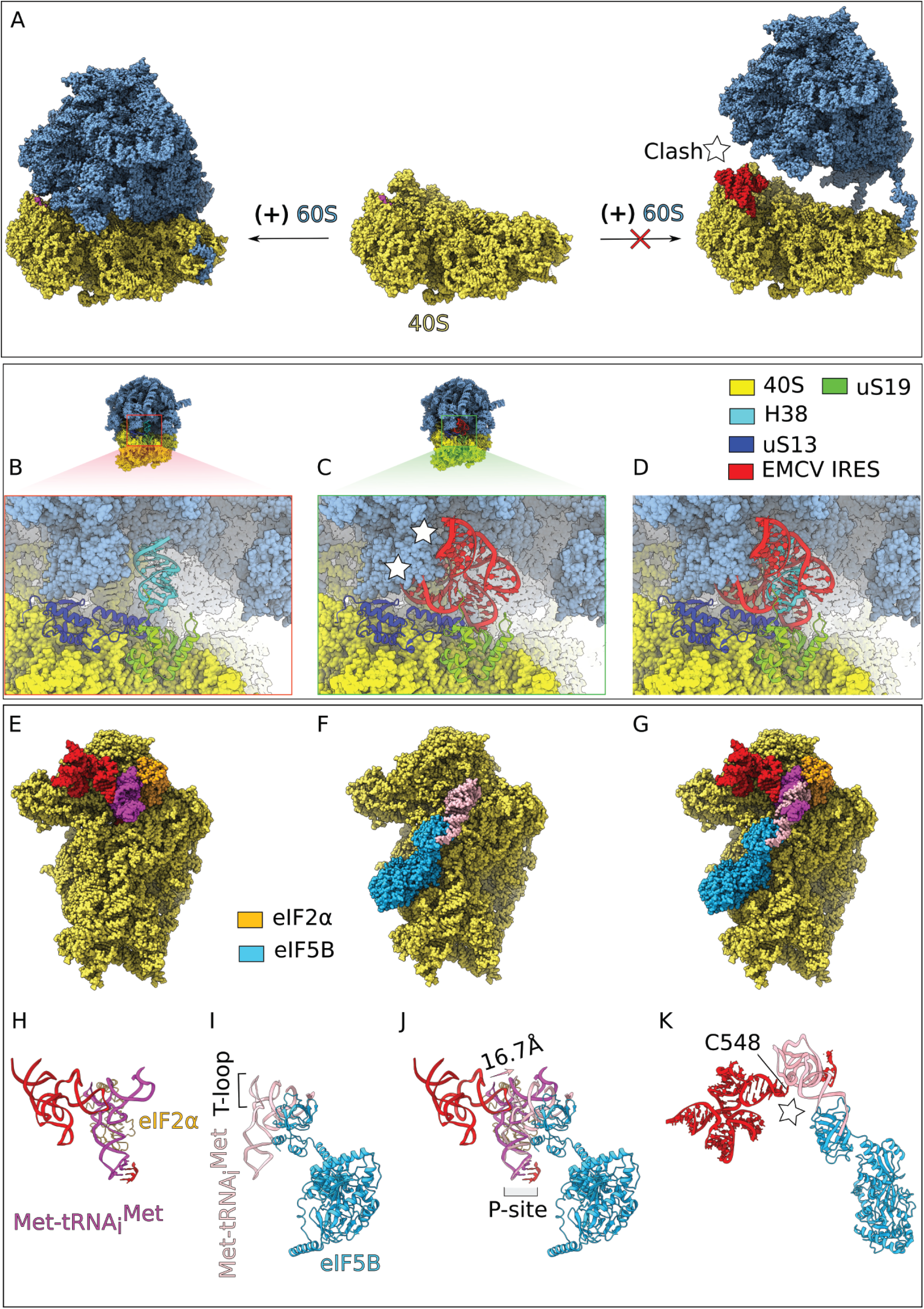
Release of the EMCV IRES from the 40S ribosomal head is necessary for joining of the 60S subunit and formation of the 80S ribosome. (**A**) On the right, formation of the 80S complex is prevented by a steric clash between the 40S-bound EMCV IRES and the 60S subunit. Removal of the EMCV IRES (red) is required to prevent a significant steric clash. (**B**) Close-up view of the 80S complex, showing that its formation is stabilized by the inter-subunit bridge B1a, which is established between H38 of the 60S subunit and uS19 of the 40S subunit. (**C**) Close-up view on a steric clash between the EMCV IRES and the 60S subunit. (**D**) The EMCV IRES appears to compete for the same binding site with uS19 that H38 occupies during 80S formation. (**E**) Interactions among the 40S subunit, the EMCV IRES, initiator tRNA (magenta), and eIF2α. (**F**) Interactions between the 40S subunit, initiator tRNA (light pink) and eIF5B after dissociation of eIF2 during subunit joining (pdb id: 7SYV). (**G**) Superimposition of 40S complexes shown in (**E**) and (**F**). (**H**) The ribosomal positioning of initiator tRNA (magenta) bound to the EMCV IRES and eIF2α in 48S initiation complexes. (**I**) The ribosomal positioning of initiator tRNA (light pink) and eIF5B in 40S ribosomal complexes in the process of subunit joining. (**J**) Superimposition of (**H**) and (**I**) showing a 16.7 Å shift in the position of tRNA, which leads to (**K**) steric interference between the tRNA T-loop and the C548 residue of the EMCV IRES tetraloop-containing subdomain after eIF2 is replaced by eIF5B, disrupting the stable tetraloop-tRNA contact.

### Mutational analysis of the central stem of domain I, the region upstream of domain I and the 3’-terminal domain L of the EMCV IRES

The importance of the structure of domain I’s long central stem has remained obscure. We investigated it by introducing disruptive/stabilizing and shortening/lengthening mutations in its predicted structural elements (Figure 6A) and assayed the activity of such mutants in 48S complex formation in the *in vitro* reconstituted system. Shortening or lengthening of the apical helix Ih1 (mutants 1 and 2) noticeably reduced 48S complex formation (Figure 6B, lanes 3-4), whereas shortening of the lower helix (mutant 3) had a lesser effect (Figure 6B, lane 5). The activity of the IRES was very tolerant to significant changes in the lower part of the central stem, which included disruption, stabilization, truncation or extension of various regions (mutants 4-9) (Figure 6C, lanes 3-8).

**Figure 6.**
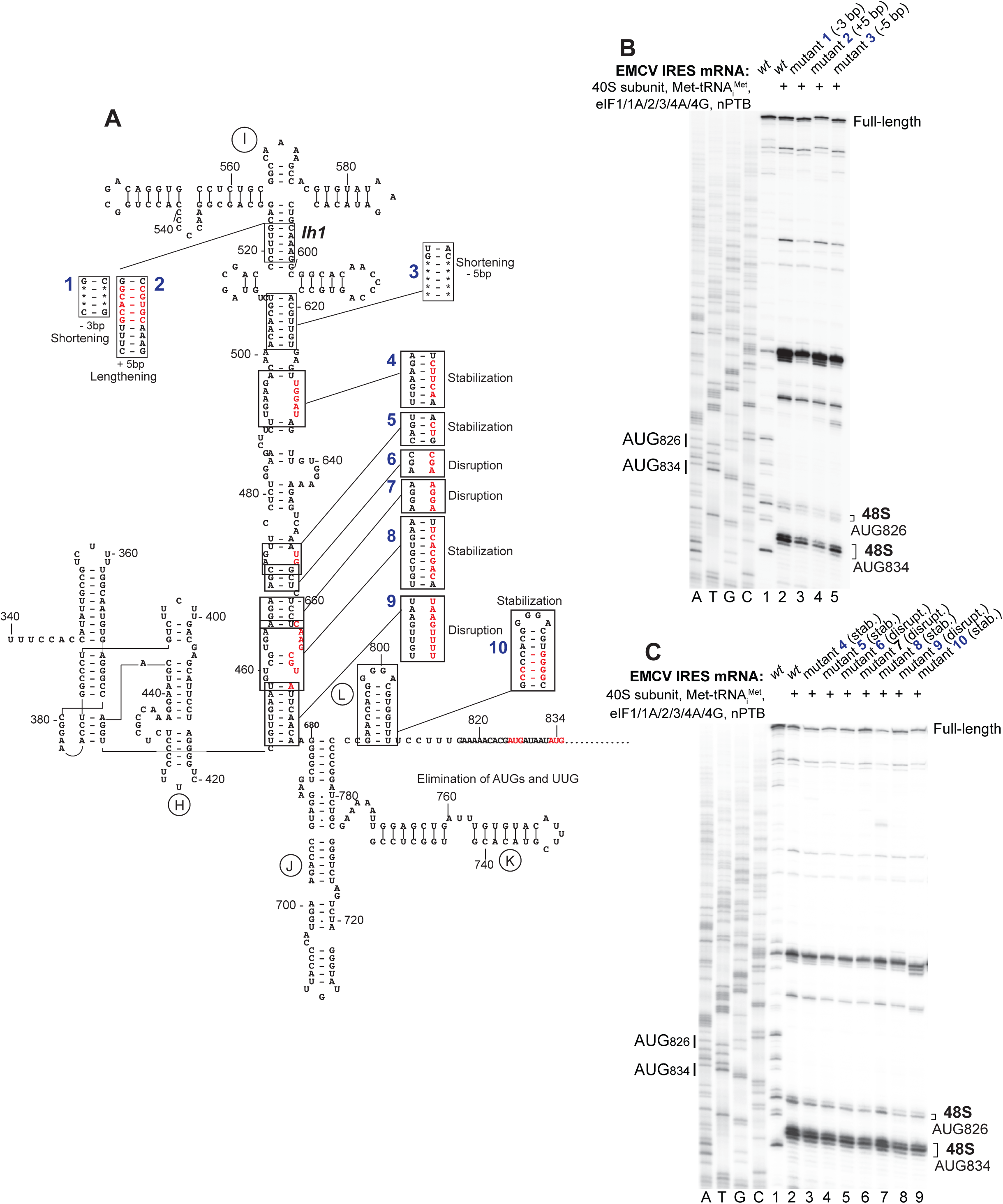
Influence of mutations in the stem of domain I on 48S complex formation on the EMCV IRES. (**A**) Secondary structure model of the EMCV IRES showing introduced mutations. (**B-C**) Toe-printing analysis of 48S complex formation on wt and mutant EMCV IRES mRNAs in the presence of individual 40S subunits, Met-tRNA_i_^Met^, nPTB and indicated eIFs. Lanes C, T, A, and G depict the wt EMCV sequence generated using the same primer. The positions of initiation codons are indicated on the left and of assembled 48S complexes on the right.

Stabilization of domain L located downstream of JK domain (mutant 10 in Figure 6A) or its entire deletion or replacement by 2 or 6 nucleotides (Figure 7A, mutants mL1-4) did not affect the activity of the IRES (Figure 6C, lane 9; Figure 7B, lanes 3-6).

**Figure 7.**
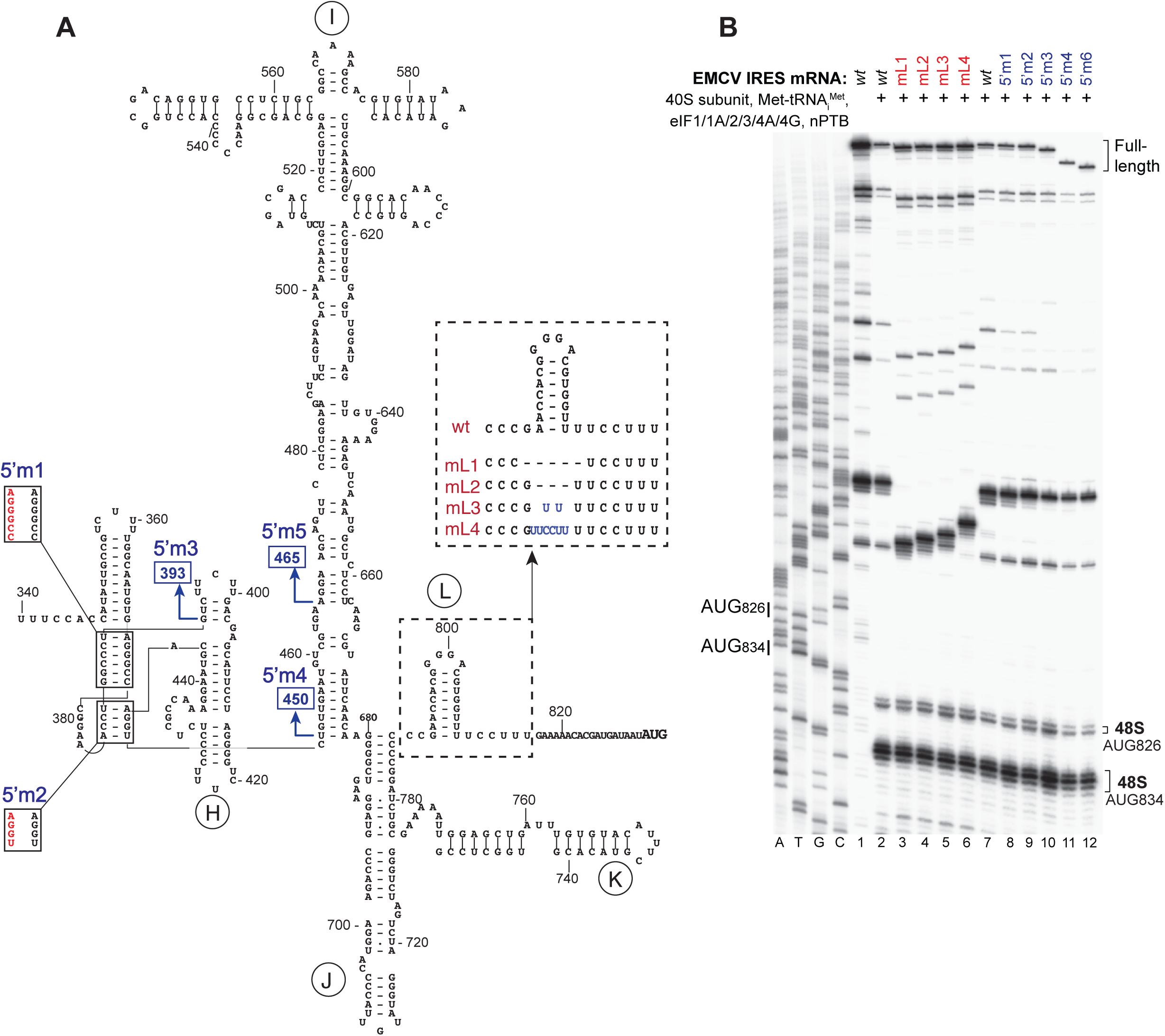
Influence of mutations in the region upstream of domain I and in domain L downstream of the JK domain on 48S complex formation on the EMCV IRES. (**A**) Secondary structure model of the EMCV IRES showing introduced mutations. (**B**) Toe-printing analysis of 48S complex formation on wt and mutant EMCV IRES mRNAs in the presence of individual 40S subunits, Met-tRNA_i_^Met^, nPTB and indicated eIFs. Lanes C, T, A, and G depict wt EMCV sequence generated using the same primer. The positions of initiation codons are indicated on the left and of assembled 48S complexes on the right.

The importance of the region upstream of domain I for IRES function is contentious (Jang and Wimmer, 1990; Duke et al., 1992), and the structure of this region is not firmly established. Modeling of the type 2 IRESs of Theiler’s murine encephalitis virus and Saffold-like cardiovirus suggest that this region forms a pseudoknot^60,61^, and that the EMCV IRES has the potential to form an analogous structure. We assayed the importance of the presence and structure of this upstream region for function using a panel of mutants (Figure 7A). Structural disruption (Figure 7A, mutants 5’m1-2) or deletion of this region (Figure 7A, mutants 5’m3-5) did not affect 48S complex formation in the *in vitro* reconstitution system (Figure 7B, lanes 8-12). Taken together, these and earlier studies^27,33,39^ identified domains I and J-K as being critical for EMCV IRES function in the *in vitro* reconstituted system, whereas upstream and downstream domains were non-essential.

## DISCUSSION

Our breakthrough cryo-EM structure of 48S complexes assembled on the EMCV IRES uncovered essential direct interactions of the apex of domain I of the IRES with the 40S subunit and initiator tRNA. Regarding the high conservation of primary and secondary structures of the apex of domain I among Type 2 IRES, and particularly of the elements directly involved in the interaction with 40S subunits and initiator tRNA (Figure S8, Table S3) as well as mutagenesis data supporting the importance of this region for the function of these IRESs^46–52^, the interactions observed for the EMCV IRES must be common for all Type 2 IRESs.

Prior to our study, direct ribosomal interaction was considered as the driving force for initiation only for HCV-, HalV- and CrPV-like IRESs, and the only known essential interaction of Type 2 IRESs involved eIF4G. Direct specific interaction of Type 2 EMCV IRES with the 40S subunit revealed in our study indicates that intimate contacts with the 40S subunit is a general common characteristic of diverse initiation mechanisms utilized by different viral IRESs. Interestingly, all of these interactions utilize regions of the 40S subunits that are used for binding of different canonical components of the translational apparatus. Thus, ribosomal binding of CrPV- and HalV-like type 6 IRESs involves A and P sites, respectively^9,10^, whereas HCV-like type 4 IRESs usurp the binding site for the structural core of eIF3^7^. In contrast to HCV-, HalV- and CrPV-like IRESs which do not use all canonical initiation components for 48S complex formation and can therefore usurp and employ their binding sites, initiation on Type 2 IRESs requires initiator tRNA and a full set of eIFs. Thus, ribosomal binding of the EMCV IRES includes interaction with uS19, which interacts with H38 (the A-site finger) of the 60S subunit to form the B1a intersubunit bridge^58,59^, a translational component that does not participate in 48S complex formation.

Although the interaction of Type 2 IRESs with individual 40S subunits has been reported, binding was rather weak (equilibrium dissociation constant (*K*_d_) of 55 ± 10 nM)^62^, compared to 1.9 ± 0.3 nM for the HCV IRES^63^ and of unclear specificity, since protection from SHAPE modification was observed in domain G (upstream of domain H) and in the nonconserved central region of domain I. Our structure shows that the interaction of the apex of domain I with the 40S subunit is supported and stabilized by the specific interaction of its GNRA loop with the initiator tRNA TψC domain. Interactions of the apex of domain I with 40S subunits and initiator tRNA in 48S ICs pose the key question of whether they can occur independently of the JK domain’s interaction with eIF4G/eIF4A and thus act as the primary interactions responsible for the ribosomal recruitment of Type 2 IRESs. Although the interaction of the JK domain with eIF4G/4A could likely aid ribosomal recruitment of Type 2 IRESs, the continuous association of the apex of domain I with initiation complexes even after dissociation of the eIF4G/4A-bound JK domain during grid preparation suggests that it is sufficiently strong and might therefore be the primary interaction responsible for the ribosomal recruitment of Type 2 IRESs.

Binding of the apex of domain I to the 40S subunit creates a steric clash with the 60S subunit indicating that the IRES must be released to allow subunit joining. Strikingly, the interaction of the IRES with initiator tRNA not only represents a contact that could facilitate ribosomal recruitment of the IRES but also provides a potential mechanism for its displacement. Thus, upon dissociation of eIF2 following eIF5-induced hydrolysis of eIF2-bound GTP and subsequent replacement of eIF2 by eIF5B, eIF5B-bound initiator tRNA undergoes a 16.7 Å shift compared to its position in eIF2-containing 48S complexes^2,3^, which would result in a steric clash between the tRNA T-loop and the C548 residue of the tetraloop-containing subdomain. This clash could disrupt the tetraloop-tRNA interaction, potentially leading to release of the apex of domain I from the 40S subunit. The observed inability of eIF5B or eIF2D to substitute for eIF2 in 48S complex formation on the EMCV IRES might also be explained by the specific unique position of initiator tRNA in eIF2-containing initiation complexes that is compatible with the interaction with the GNRA loop of the EMCV IRES.

Thus, Type 2 IRESs contain two major elements: the apical region of domain I which binds the 40S subunit and tRNA, and the J-K domain which binds eIF4G/eIF4A. Whereas the primary role of the former would be consistent with facilitating attachment of the IRES to 43S preinitiation complexes, the primary role of the latter is likely to promote loading of the region around the initiation codon into the mRNA-binding channel, which is supported by the observation that eIF4A/4G induce ATP-dependent conformational changes in this region of Type 2 IRESs^39^. The fact that the activity of the EMCV IRES was very tolerant to significant changes in the lower part of the central stem of domain I (disruption, stabilization, truncation or extension of various regions) (Figure 6) suggests that its role might be merely as a connector between the two major functional elements. Consistent with this idea, the lower part of the central stem of domain I is the least conserved region in Type 2 IRESs^64,65^. The region upstream of domain I as well as domain L located downstream of J-K domain also were not essential for the activity of the EMCV IRES in the in vitro reconstituted system (Figure 7), emphasizing further the exceptional role of two major elements: the apical region of domain I and the J-K domain.

The key remaining question in the initiation on Type 2 IRESs is the exact mechanism of action of eIF4G/4A and the role in it of their specific association with the JK domain. To avoid potential artifacts, we did not perform chemical cross-linking of the assembled 48S complexes used for cryo-EM studies, even though cross-linking stabilizes the association eIF3, eIF4A and eIF4G with ribosomal complexes^54,66^. As expected, the structure of uncross-linked 48S complexes did not reveal the position of the eIF4G/4A-bound JK domain. Initiation on the EMCV IRES does not require the portion of eIF4G that is responsible for its interaction with eIF3^38,53^, which poses the following questions: (i) does the JK domain play the role in positioning of the eIF4G/eIF4A/JK domain complex by specific interactions with e.g. the 40S subunit or eIF3? and (ii) is the position of eIF4G/eIF4A in initiation complexes assembled on the EMCV IRES the same as in canonical initiation complexes or does the JK domain usurp eIF4G/eIF4A and displace them from their conventional binding site analogously to how the HCV IRES binds eIF3 and displaces it from its position on the 40S subunit^7^? Thus, to elucidate the molecular mechanism of action of eIF4G/4A during initiation on Type 2 IRESs, future studies should focus on obtaining structures of 48S complexes showing the eIF4G/eIF4A-bound JK domain.

## RESOURCE AVAILABILITY

### Lead contact

For any inquiries, contact Tatyana Pestova.

### Materials availability

Plasmids generated for this study are available by request to the lead contact.

### Data and code availability

The cryo-EM density maps have been deposited in the Electron Microscopy Data Bank (EMDB) under accession numbers EMD-40769, EMD-40770, EMD-40771, EMD-40772, EMD-40773, and EMD-40774. The corresponding atomic model for EMD-40774 has been deposited in the Protein Data Bank (PDB) under accession code 8SUP.

## ACKNOWLEDGMENTS

We thank Swastik De and Jeevan GC for providing valuable suggestions. All data were collected at the Cryo-EM core of Columbia University. We thank Robert A. Grassucci, Zhening Zhang, and Yen-Hong Kao for their help with the cryo-EM data collection. This work was supported by NIH grants R01GM29169 and R35GM139453 to JF, R35GM122602 to TVP, and R01GM097014 and R21AI188505 to CUTH.

## AUTHOR CONTRIBUTIONS

The project was conceived, and experiments designed, by C.H., J.F., and T.P. I.A. prepared the biological samples for cryo-EM studies. Y.A. performed biochemical experiments. H.F. prepared cryo-EM grids and collected data. Z.P.B. processed cryo-EM data, performed structure determination, and built and refined atomic models. S.B. collected additional cryo-EM data, processed the merged dataset, built the final set of atomic models, and prepared figures. Z.B., S.B., C.H. and T.P. analyzed and interpreted the models in terms of viral IRES-launched translation initiation. The manuscript was written by T.P., C.H., J.F., S.B., and Z.P.B with the input from I.A and Y.A.

## MATERIALS AND METHODS

### Construction of plasmids

The transcription vector for human tRNA_i_^Met^ has been described^68^. The EMCV transcription vectors T7-EMCV(373-1656)wt_pUC57 and T7-EMCV(373-987)wt_pUC57, containing a T7 promoter followed by EMCV nt. 373-1656 or EMCV nt. 373-987 (GenBank: M81861.1) inserted into pUC57, and mutants derived from these plasmids were made by GenScript Corp. (Piscataway, NJ) and by Synbio Technologies (Monmouth Junction, NJ), respectively. The EMCV plasmids were linearized by EcoRI, and mRNAs were transcribed using T7 RNA polymerase (Thermo Scientific). The plasmid pTZ18R-EMCV(315-1160) containing a T7 promoter followed by EMCV nt. 315-1160, with ATG826-828ATT and TGC861-863AGT substitutions in the EMCV sequence to eliminate the weakly used AUG826 (and thus to improve the homogeneity of the 48S complexes) and to add an Spe1 site (for restriction, to yield mRNA that terminates at nt. 862) was used for generating mRNA employed in assembly of 48S complexes for cryo-EM studies, directed hydroxyl radical cleavage experiments and enzymatic footprinting.

Vectors for expression of His_6_-tagged wild type eIF1 and eIF1A^28^, eIF4A and eIF4B^26^, eIF4G1(653–1599)^27^, nPTB^60^, *Escherichia coli* methionyl tRNA synthetase^69^, eIF2D^13^, and of eIF1A proteins containing unique surface-exposed cysteines^55^ have been described.

### Purification of ribosomal subunits, initiation factors and aminoacyl-tRNA synthetases

Native mammalian 40S subunits, eIF2, eIF3, eIF4F and eIF5B were purified from rabbit reticulocyte lysate (RRL) (Green Hectares, Oregon, WI) as described^70^. Human recombinant eIF1, eIF1A, eIF4A, eIF4B, eIF4G_653-1599_, nPTB, eIF2D and *E. coli* methionyl-tRNA synthetase were expressed in *E. coli* BL21(DE3) (Invitrogen) and purified as described^13,26–28,38,69^.

Native total calf liver tRNA (Promega) and *in vitro* transcribed tRNA_i_^Met^ were aminoacylated using recombinant *E. coli* methionyl-tRNA synthetase^69,70^.

### Assembly of 48S complexes on the EMCV IRES for cryo-EM studies

48S complexes were assembled by incubating 9 pmol EMCV(ATG826-828ATT/TGC861-863AGT) mRNA with 3 pmol 40S subunits, 12 pmol eIF2, 6 pmol eIF3, 6 pmol eIF4F, 12 pmol eIF4A, 10 pmol eIF4B, 15 pmol eIF1, 15 pmol eIF1A, 10 pmol nPTB and 5 pmol native Met-tRNA_i_^Met^ in 40 μl buffer A (20 mM Tris pH 7.5, 100 mM KCl, 2.5 mM MgCl_2_, 2 mM DTT and 0.25 mM spermidine) supplemented with 1mM ATP and 0.4 mM GTP for 15 min at 37°C.

### Preparation of cryo-EM grid and image acquisition

Copper Quantifoil grids R0.6/1.0 with mesh size 300 (Quantifoil Micro Tools GmbH) were subjected to glow discharge with air for 30 s using a PELCO easiGlow cleaning system set to a plasma current of 15 mA, to make the carbon film surface negatively charged, which allows buffer solutions to spread easily. These grids were immediately placed into the Vitrobot Mark IV (Thermo Fisher Scientific, Carlsbad, CA) maintained at 4°C and 98% humidity. 3 μl of the above-mentioned sample solution was applied onto the grid and blotted for 4 s with blot force 3. The sample containing grid was plunged into liquid ethane and then carefully transferred to liquid nitrogen. Grids were clipped and screened to confirm sample distribution and ice thickness using a Tecnai F20 electron microscope (Thermo Fisher Scientific) equipped with a field emission gun (FEG) operating at 200 kV and a K3 Summit direct electron detector (Gatan, Inc, Pleasanton, CA).

After screening the best grids are selected and further data was collected on a Tecnai Polara F30 and Titan Krios (Thermo Fisher Scientific) microscopes operated at 300 KeV equipped with a K3 direct detector (Gatan). Movies were collected at a pixel size of 0.95 Å/pixel for Tecnai Polara F30 and 0.83 Å/pixel for Krios. The defocus range was set from -1 to -2.5 μm. A total of 7733 micrographs were collected. 40S ribosomal subunits showed a preferred orientation, so, portions of the data were collected with a 35° stage tilt.

### Image processing

A workflow of the data processing for the cryo-EM image was shown in Figure S2. Drift, gain correction, and dose weighting were performed using MotionCor2^71^ with a local patch correction with 7 × 5 patches. The contrast transfer function (CTF) of each micrograph was estimated using the CTFFIND4^72^. Particle picking was performed using Topaz^73^. Good particles were selected by 2D classification and trained using 20,000 particles in Relion4.0^74–77^. Autopicking using the trained topaz model yielded 835,875 particles. Particles picked by Topaz were subjected to four rounds of 2D classification for a further selection of good particles, which yielded 688,588 particles. All particles were pooled together and used for initial model generation followed by 3D auto-refinement, applying C1 symmetry in Relion4.0. CTF refinements were done to correct for magnification anisotropy, fourth-order aberrations, per-particle defocus, and per-particle astigmatism, followed by another 3D auto-refinement. The final resolution after refinement was 3.2 Å (Figure S3). Then 3D classification was performed without alignment, using the angular information from the previous refinement step. Four distinct classes—Class I (134,626 particles), Class II (98,748 particles), Class III (89,242 particles), and Class IV (72,526 particles)—were identified, all exhibiting clear density for the 40S ribosomal subunit, eIF1A and the IRES bound to the inter-subunit face of the 40S head. Classes II-IV also contained density for eIF1, whereas not fully resolved Class I contained additional density for eIF2, Met-tRNA_i_^Met^, and blurry density for eIF3. Additionally, Classes II–IV differ from one another in the extent of 40S head rotation. Focused classification of the Class I population yielded a refined subclass, termed Class Ia, comprising 59,153 particles with well-resolved density for the 40S, eIF2, eIF1A, Met-tRNA_i_^Met^, and the IRES. We further did multiple rounds of focused refinement on the 40S head along with associated EMCV mRNA and Met-tRNA_i_^Met^ regions, and this yielded a 3.1 Å reconstructed map. In addition, series of focused 3D classification, 3D Flexible Refinement (https://guide.cryosparc.com/processing-data/tutorials-and-case-studies/tutorial-3d-flex-mesh-preparation), Blush regularization^78^ on EMCV mRNA, 40S-head, and Met-tRNA_i_^Met^ regions and followed by relion_reconstruct with Ewald sphere correction were performed to resolve the tetraloop-tRNA-40S-head interacting region. All raw FSC curves representing the resolution estimations are shown in Figure S3.

### Model building and refinement

For starting models we used available models for the 48S IC (PDB: 7SYR)^2^ and eIF3 (PDB: 5A5T)^79^. Initial rigid body model fitting was performed using UCSF Chimera v1.1^80^ with additional manual flexible fitting in Coot^81^. All models were further flexibly fitted using Phenix geometry minimization and multiple rounds of PHENIX real-space refinement^82,83^. The initial model for the EMCV domain-I (residues 518-600) was built by using 3dRNA^84^ restraints based on the connectivity of the bases from the 2D structure).

### Determining EMCV mRNA sequence nucleotide identity

The EMCV genome (GenBank: M81861.1; https://www.ncbi.nlm.nih.gov/nuccore/M81861.1) contains an IRES sequence of 441 nucleotides in length covering bases 396UUC…AUG836. The density of IRES visible in our map is approximately 80-90 nucleotides and so we had to determine which region of the complete IRES was visible. To determine which region of the IRES is visible in our density maps (Figure S4B), we first identified the highest resolution portion, a loop bound into the pocket between alpha helix 2-3 of ribosomal protein uS13 and the beta-sheet of uS19. Here we could identify sequential purine (R) and pyrimidine (Y) bases (Figure S4C). This region of the IRES contained 14 nucleotides where their identity could be determined giving a putative sequence of YRYRRYNNNRNNYY. Since the directionality of the sequence cannot be determined at this resolution, we performed the following search in both the 5ʹ to 3ʹ and 3ʹ to 5ʹ directions (e.g., (5ʹ)YRYRRYNNNRNNYY(3ʹ) and (3ʹ)YRYRRYNNNRNNYY(5ʹ)). As will be detailed below, the 5ʹ to 3ʹ case proved to be correct with the observed density corresponding to 561TGCGGCCAAAAGCC574, so it will be explained first, but the 3ʹ to 5ʹ details will also follow.

### The (5ʹ)RNRR(3ʹ) direction

The leading RYRR region contained the most consecutive purines, which given their size could be most confidently identified, and was used as an initial search term to find candidate matches within the IRES gene. Using the minimal search term of RNRR we identified 40 locations (excluding overlapping regions) that were present across all domains of the IRES. There were four matches in domain H (402ACGA405; 414AGGG417; 435AAAG438; 443GCAA446), 24 in domain I (462GTGA465; 466AGGA469; 471GCAG474; 482GGAA485; 493GAAG496; 497ACAA500; 501ACAA504; 510GTAG513; 523GCAG526; 527GCAG530; 532GGAA535; 547GCGA550; 562GCGG565; 568AAAA571; 581ATAA584; 594GCAA597; 600GCGG603; 605ACAA608; 624GTGA627; 633ATAG636; 639GTGG642; 643AAAG646; 653ATGG656; 676ACAA679), five in domain J (680GGGG683; 686GAAG689; 697AGAA700; 713ATGG717; 728GGGG731), two in domain K (764GAGG767; 770AAAA773), two in domain L (795ACGG798; 803GTGG806), and three in the unstructured region containing the start codon (816GAAA819; 823ACGA826; 829ATAA832). Narrowing down the potential matches was done in two ways, either by correlating with the known secondary structure of the EMCV IRES, or by increasing the search term to include all confidently identified nucleotides. Both methods independently return the same result.

### Using the IRES secondary structure

The visible sequence (YRYLOOPRRYNNNRNNYYLOOP) contains a loop and so using the EMCV IRES secondary structure we can eliminate matches that occur within helices or regions that contain loops that are too short or too long to match the observed density. The loop region contains 11 nucleotides (Figure S4C). So we examined the EMCV secondary structure and found 10 candidates that were within a loop region: one in domain H (414AGGG417), eight in domain I (510GTAG513; 532GGAA535; 547GCGA550; 562GCGG565; 568AAAA571; 581ATAA584; 600GCGG603; 605ACAA608), and one in domain L (795ACGG798). All matches in the unstructured region containing the start codon were excluded. Of the 10 candidate locations, one would require a loop that is too long, 17 nucleotides for 600GCGG603, and the other nine loops that are too short, seven nucleotides for 532GGAA535 and 605ACAA608, six nucleotides for 795ACGG798, four nucleotides for 510GTAG513, three nucleotides for 581ATAA584, and two nucleotides for 547GCGA550 and 568AAAA571. None of these were appropriate even after considering ambiguity in the number of observed nucleotides as the longest would require an error of (+6) nucleotides (600GCGG603) and the next shortest an error of (-4) nucleotides (532GGAA535; 605ACAA608), something that is not supported by the observed data. The remaining two locations that were in a region of the IRES that could support a loop of 11 nucleotides were 414AGGG417 in domain H and 562GCGG565 in domain Id. Distinguishing between these two locations can be completed using different methods. First, the global arrangement of the IRES suggests a cloverleaf conformation, something that is absent in domain H but present in domain Id (Figure S1). Alternatively, considering the second nucleotide and assuming it’s accurately identical to a pyrimidine (e.g., RYRR) this also eliminates domain H (414AGGG417) and leaves only domain Id (562GCGG565).

### Using a longer search term

Disregarding the constraints given by the secondary structure we can also identify domain Id as the location of the IRES density by increasing the length of the search term. We used every combination from RNRR to the full set of visible nucleotides (all terms and results given in Table S2) and found that YRYRRYNNNRNNYY corresponds with 561TGCGGCCAAAAGCC574 in domain Id. No other region in the IRES had a positive result for the complete search term.

### The (3ʹ)RNRR(5ʹ) direction

Given the resolution, it is not possible to determine the directionality of the high-resolution loop region and so we also searched in the 3ʹ to 5ʹ direction. Searching for RRNR we identified 44 locations (excluding overlapping regions) that were present across all domains of the IRES. There were six matches in domain H (401GACG404; 405AGCA408; 414AGGG417; 435AAAG438; 440AATG443; 445AAGG448), 23 in domain I (456AATG459; 464GAAG467; 468GAAG471; 482GGAA485; 493GAAG496; 500AACA503; 512AGCG515; 526GGCA529; 532GGAA535; 546GGCG549; 553GGTG556; 568AAAA571; 583AAGA586; 596AAAG599; 602GGCA605; 613AGTG616; 632GATA635; 641GGAA644; 645AGAG648; 652AATG655; 666AGCG669; 675AACA678; 679AGGG682), six in domain J (686GAAG689; 690GATG693; 697AGAA700; 701GGTA704; 716GGGA719; 728GGGG731), four in domain K (736GGTG739; 764GAGG767; 770AAAA773; 774AACG777), one in domain L (797GGGG800), and four in the unstructured region containing the start codon (816GAAA819; 820AACA823; 825GATG828; 831AATA834). Narrowing down the potential matches was done in two ways, either by correlating with the known secondary structure of the EMCV IRES, or by increasing the search term to include all confidently identified nucleotides. Both methods independently eliminated all results and therefore strongly suggested that the correct directionality is not (5ʹ)RRNR(3ʹ) but the (5ʹ)RNRR(3ʹ) described in detail above.

### RRNR: Using the IRES secondary structure

The visible sequence (LOOPYYNNRNNNYRRLOOPYRY) contains a loop and so using the EMCV IRES secondary structure we can eliminate matches that occur within helices or regions that contain loops that are too short or too long to match the observed density. The loop region contains 11 nucleotides (Figure S4C) and so we examined the EMCV secondary structure and found 5 candidates that were within a loop region: three in domain I (512AGCG515; 553GGTG556; 568AAAA571), one in domain J (716GGGA719), and one in domain L (797GGGG800). These would require loops of length 13 (716GGGA719), 12 (553GGTG556), and two (512AGCG515; 568AAAA571; 797GGGG800). The region in domain Ic (553GGTG556) that could be linked with a loop of length 12 (so only an error of one nucleotide) bares closer examination.

### RRNR: using a longer search term

Increasing the length of the search term eliminates all matches (all terms and results given in Table S2) so that the maximum search YYNNRNNNYRRYRY returns zero matches.

### Model building IRES region 518CC..GG600

Once we had determined the identity of the 561TGCGGCCAAAAGCC574 region we were able to build domains Id, Ic, Id, and Id spanning 83 nucleotides (518CC..GG600) and accounting for 17% of the IRES.

### Analysis of the activity of EMCV IRES mutants in 48S complex formation in the *in vitro* reconstituted system

48S complexes were assembled by incubating 0.3 pmol of *wt* or mutant EMCV IRES mRNAs with 1 pmol 40S subunits, 1.5 pmol native or *in vitro* transcribed Met-tRNA_i_^Met^ and indicated combinations of 5 pmol eIF2, 2 pmol eIF3, 8 pmol eIF4G_653-1599_, 8 pmol eIF4A, 3 pmol eIF4B, 10 pmol eIF1, 10 pmol eIF1A, 5 pmol nPTB, 8 pmol eIF5B and 8 pmol eIF2D in 20 μl of buffer A (20 mM Tris pH 7.5, 100 mM KCl, 2.5 mM MgCl_2_, 2 mM DTT and 0.25 mM spermidine) supplemented with 1mM ATP and 0.4 mM GTP for 15 min at 37°C. Formation of 48S complexes was monitored by toe-printing using AMV reverse transcriptase (Promega) and ^32^P-labeled primer (5’-CGGTATTGTAGAGCAG-3’) complementary to EMCV nt. 901-916 or (5’-GCAGGTAAAATCCATTACGG-3’) complementary to EMCV nt. 914-933^70^. Radiolabeled cDNAs were phenol-extracted, ethanol-precipitated, resolved on 6% polyacrylamide gel and analyzed by Phosphoimager.

### Directed hydroxyl radical cleavage

eIF1A Cys mutant proteins were derivatized with Fe(II)-BABE by incubating 3000 pmol eIF1A with 1-mM Fe(II)-BABE in 100 ml buffer B (80 mM HEPES, 300 mM KCl, 10% glycerol) for 30 min at 37°C as described^55^. Derivatized proteins were separated from unincorporated reagent by buffer exchange on Amicon 10k filter units and stored at -80°C.

48S/[Fe(II)-BABE]-eIF1A complexes were formed by incubating 0.6 pmol EMCV(ATG826-828ATT/TGC861-863AGT) mRNA with 2 pmol 40S subunits and 20 pmol [Fe(II)-BABE]-eIF1A, 3 pmol Met-tRNA_i_^Met^, 10 pmol eIF2, 4 pmol eIF3, 20 pmol eIF1, 16 pmol eIF4G_653-1599_, 16 pmol eIF4A, 6 pmol eIF4B and 10 pmol nPTB for 15 min at 37°C in 40 µL buffer A with reduced DTT (20 mM Tris, pH 7.5, 0.25 mM spermidine, 2.5 mM MgCl_2,_ 0.1 mM DTT) supplemented with 1 mM ATP, 0.5 mM GTP and 30 U RNAse inhibitor for 15 min at 37°C. To generate hydroxyl radicals, reaction mixtures are incubated on ice for 10 min and then supplemented with 0.06% H_2_O_2_ and 5 mM ascorbic acid and incubated on ice for 10 min^55^. Reactions were quenched by adding 20 mM thiourea. mRNA was then phenol-extracted, ethanol-precipitated and analyzed by primer extension using AMV reverse transcriptase and a γ^32^P-end-labelled primer (5’-GCCCCTTGTTGAATACGCTT-3’) complementary to EMCV nt. 665-684. Resulting radiolabeled cDNAs were also phenol-extracted, ethanol-precipitated, resolved on 6% polyacrylamide gel and analyzed by Phosphoimager.

### Enzymatic footprinting

48S and RNP complexes were assembled by incubating 0.6 pmol EMCV(ATG826-828ATT/TGC861-863AGT) mRNA with various combinations of 2 pmol 40S subunits, 3 pmol Met-tRNA_i_^Met^, 10 pmol eIF2, 4 pmol eIF3, 20 pmol eIF1, 20 pmol eIF1A, 16 pmol eIF4G_653-1599_, 16 pmol eIF4A, 6 pmol eIF4B and 10 pmol nPTB for 15 min at 37°C in 40 µL buffer A supplemented with 1 mM ATP, 0.5 mM GTP and 30 U RNAse inhibitor. Assembled 48S complexes were either left in the reaction mixture or purified by sucrose density gradient centrifugation essentially as described^85^. Thus, 48S complexes are assembled as described above in scaled-up 300 µL reaction mixtures and analyzed by centrifugation through 10–30% sucrose density gradients (SDGs) prepared in buffer A in a Beckman SW55 rotor at 53 000 rpm for 105 min at 4°C. The optical density of fractionated gradients was measured at 260 nm.

Individual mRNA and assembled 48S and RNP complexes were then enzymatically digested with RNase T1 as described^86^. mRNA was phenol-extracted, ethanol-precipitated, and analyzed by primer extension using AMV reverse transcriptase and γ^32^P-end-labelled primers (5’-GCAAGTCTCTTGTTCCATGG-3’) complementary to EMCV nt. 863-844 and (5’-GCCCCTTGTTGAATACGCTT-3’) complementary to EMCV nt. 665-684. The resulting radiolabeled cDNAs were also phenol-extracted, ethanol-precipitated, resolved on 6% polyacrylamide gel and analyzed by Phosphoimager.

### Determination of a consensus structure for the apical region of domain I

RNA sequences corresponding to the apex of domain I of type 2 IRESs from members of 12 genera of *Picornaviridae* and from several unclassified picornaviruses (Supplementary Table S3) were aligned using Clustal Omega. The resulting aligned sequences (41.3 – 97.5% nucleotide sequence identity) were analyzed using R-Scape^87^ to identify statistically significant nucleotide sequence covariation (*E*-value: 0.05) and power. Cacofold^88^ was then used to build a consensus structure.

