## supplemental figures and tables for "The mechanism of ribosomal recruitment during translation initiation on Type 2 IRESs"

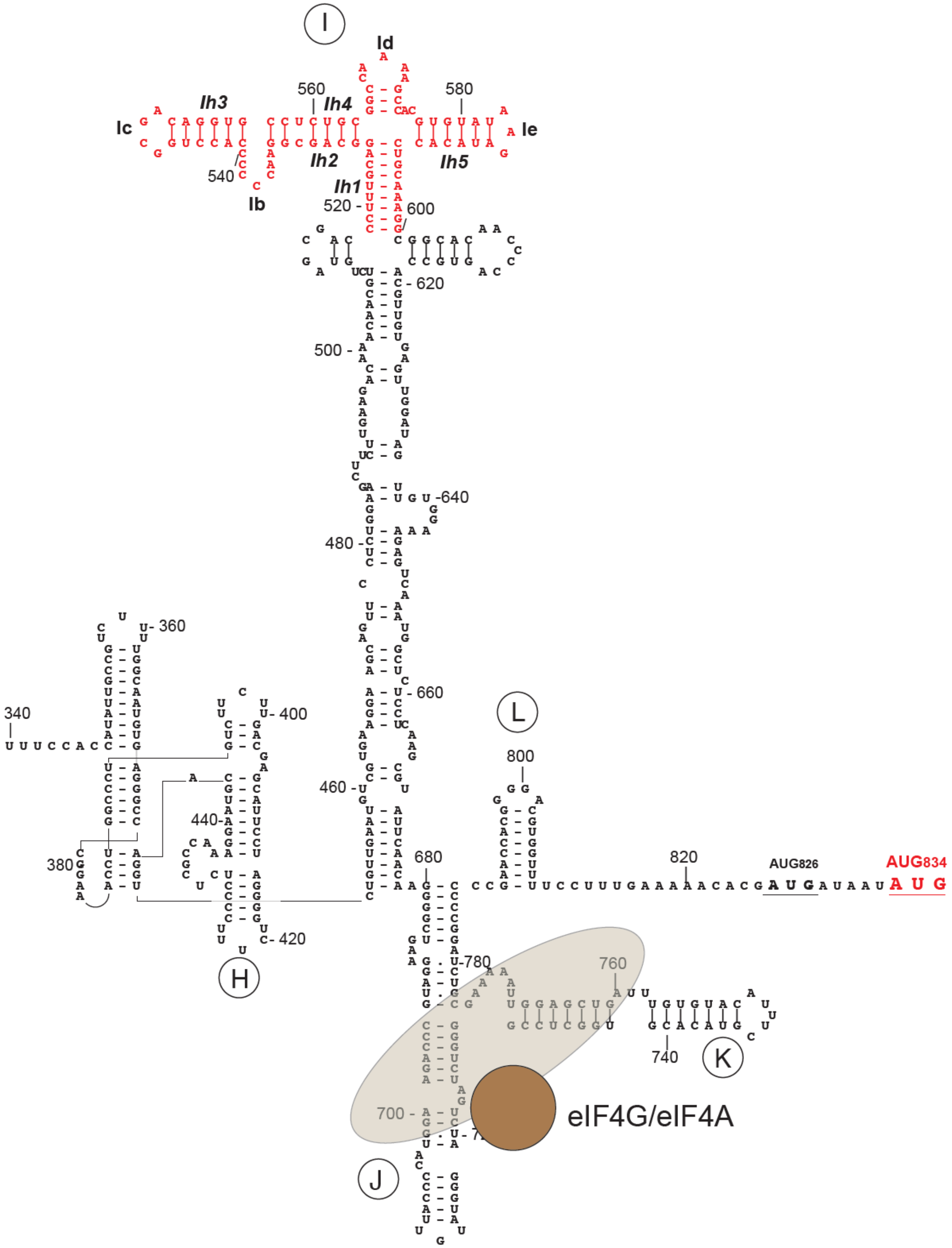


**Figure S1. The EMCV IRES secondary structure model featuring individual domains, the region corresponding to cryo-EM density (red), and specific binding of eIF4A/4G to the JK domain.**

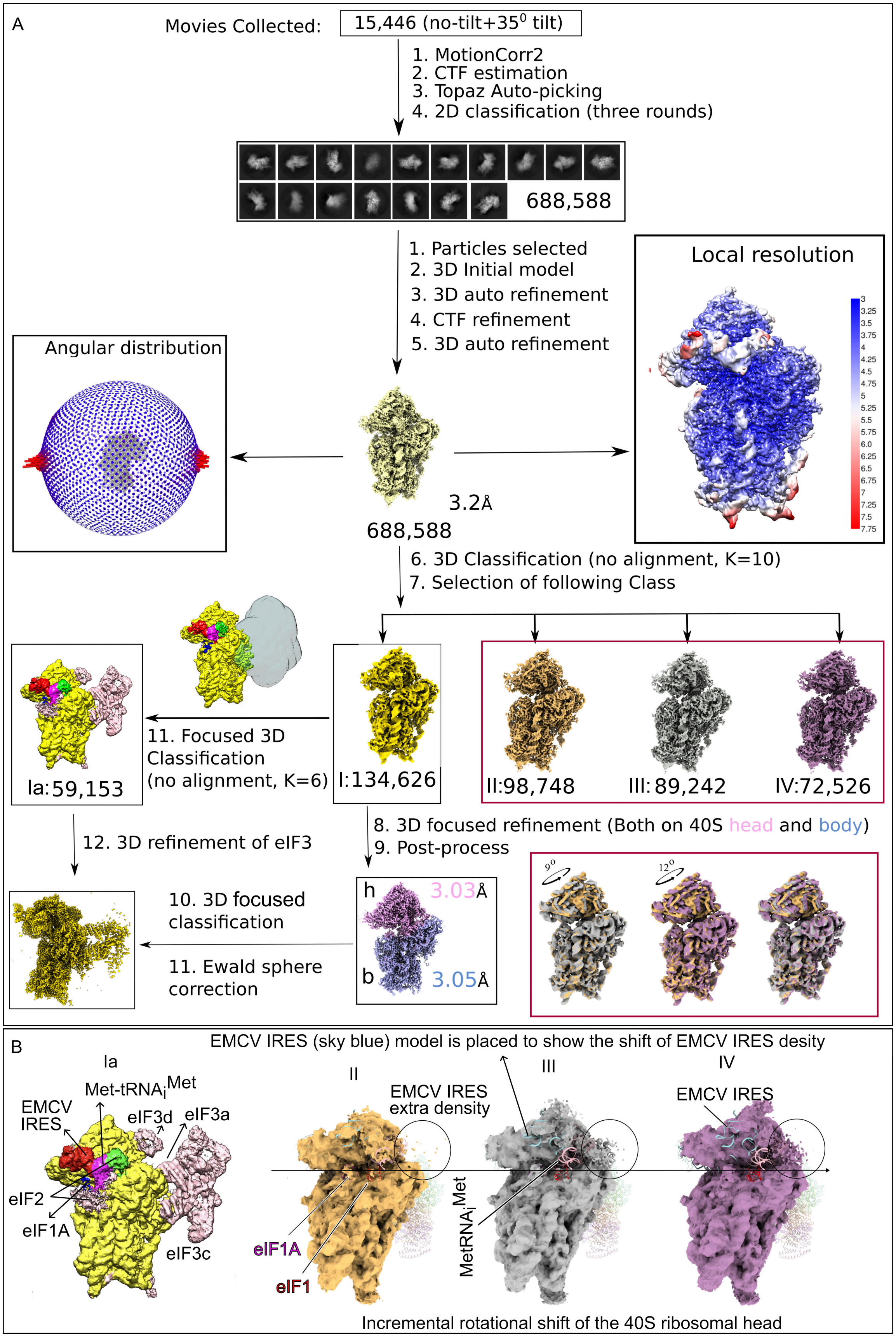


**Figure S2. Cryo-EM Data Processing Workflow and Structural Characterization of EMCV IRES-40S Initiation Complexes.**

(A) Data processing pipeline for structure determination of EMCV IRES-initiated 40S complexes. (B) Class Ia (left): 48S initiation complex showing Met-tRNA$_i^{Met}$, eIF2, eIF1A, eIF3 (eIF3a/c/d subunits are indicated), and EMCV IRES density bound at the 40S subunit head. Classes II–IV: A series of reconstructions capturing progressive 40S head rotation, accompanied by repositioning of the EMCV IRES. Fragmented extra density attributed to the IRES is highlighted (circled), and a sky-blue model of the EMCV IRES is overlaid to illustrate its positional shift. Gaussian filtering reveals this density remains connected to the IRES across all states. The horizontal arrow from left to right denotes the direction of progressive rotational movement of the 40S head, representing a continuum of structural intermediates.

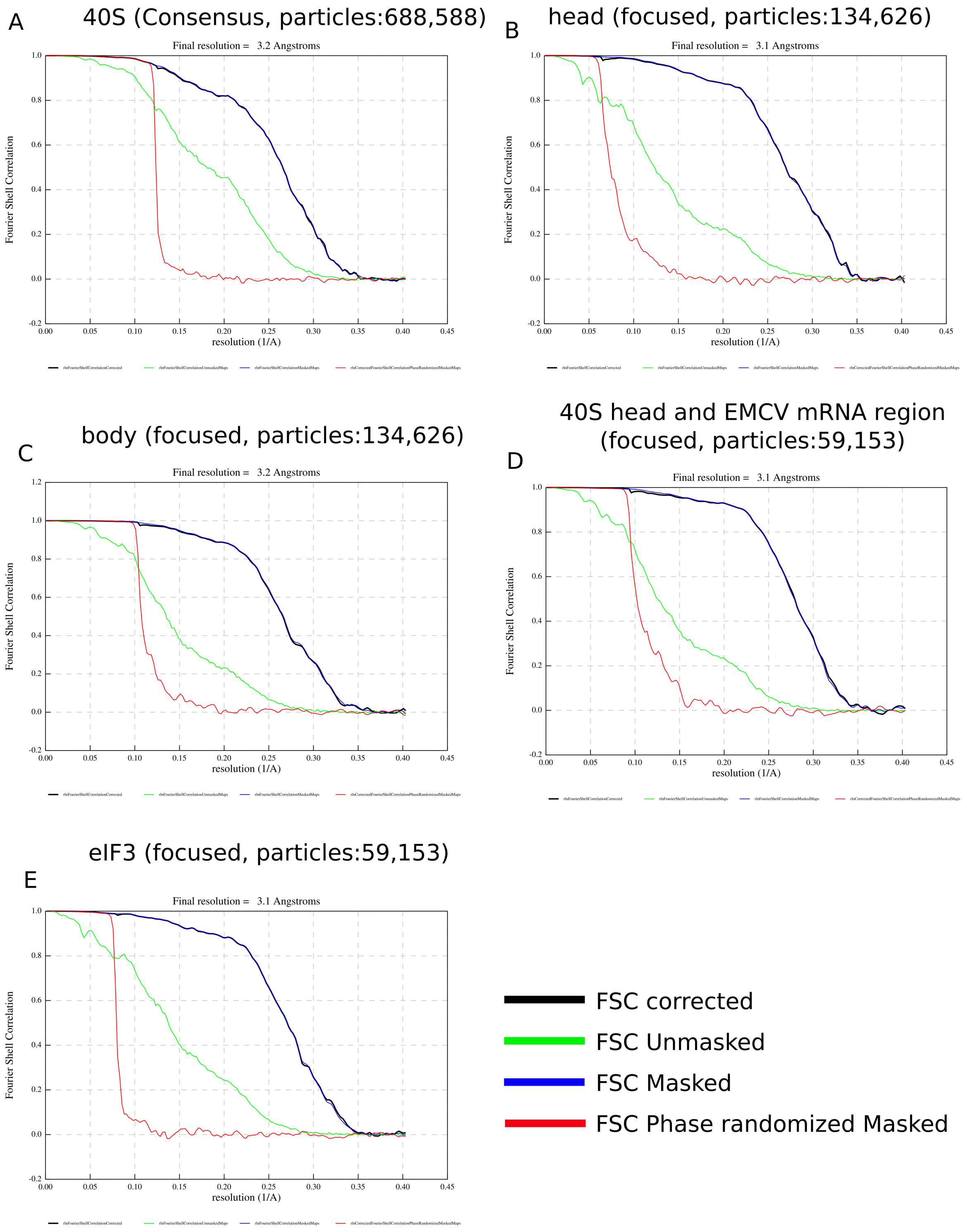


**Figure S3. Fourier Shell Correlation (FSC) curves for consensus and focused cryo-EM reconstructions.**

(A) FSC curves for the global consensus reconstruction of the 40S subunit using 688,588 particles, yielding a final resolution of 3.2 Å. (B) Focused refinement on the 40S head from 134,626 particles achieved a resolution of 3.1 Å. (C) Focused refinement on the 40S body from the same particle subset reached a resolution of 3.2 Å. (D) Refinement focused on the 40S head and EMCV mRNA region from Class Ia (59,153 particles) resulted in a 3.1 Å resolution map. (E) Focused refinement on eIF3 from Class Ia (59,153 particles) also yielded a final resolution of 3.1 Å. FSC curves are shown for corrected (black), unmasked (green), masked (blue), and phase-randomized masked (red) data, with the 0.143 criterion used to determine final resolutions.

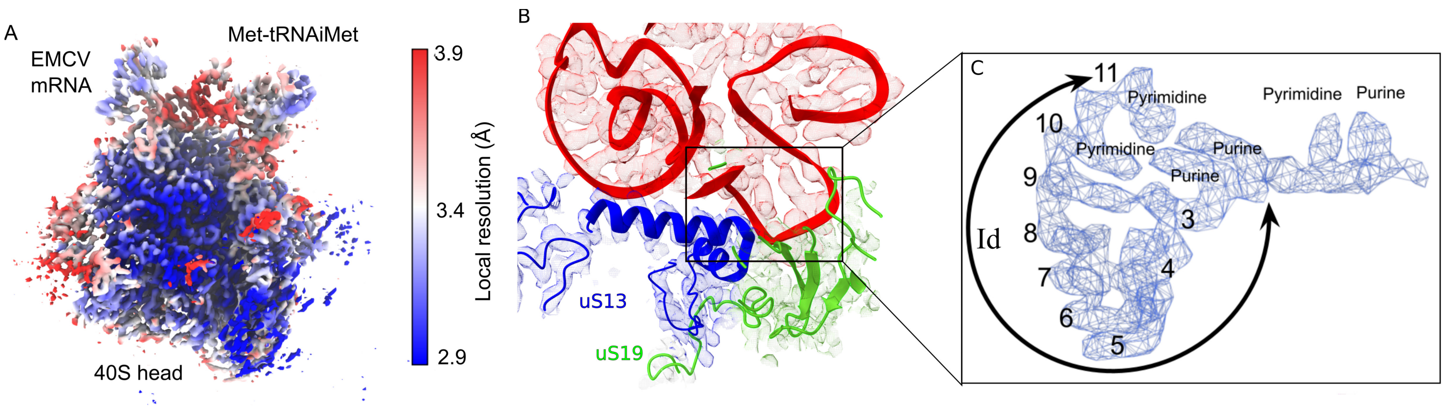


**Figure S4. Local resolution estimation and assignment of nucleotides of EMCV mRNA**.

(A) The local resolution estimation of the 40S head part along with the EMCV IRES and Met-tRNA$_i^{Met}$. (B-C) The zoomed view of the Id subdomain of the EMCV IRES (in mesh) showing the loop formation through base pairing among RNA bases. The models fitted within densities of the EMCV IRES (red), uS19 (green), and uS13 (blue) are showing the relative position of Id on the 40S head.

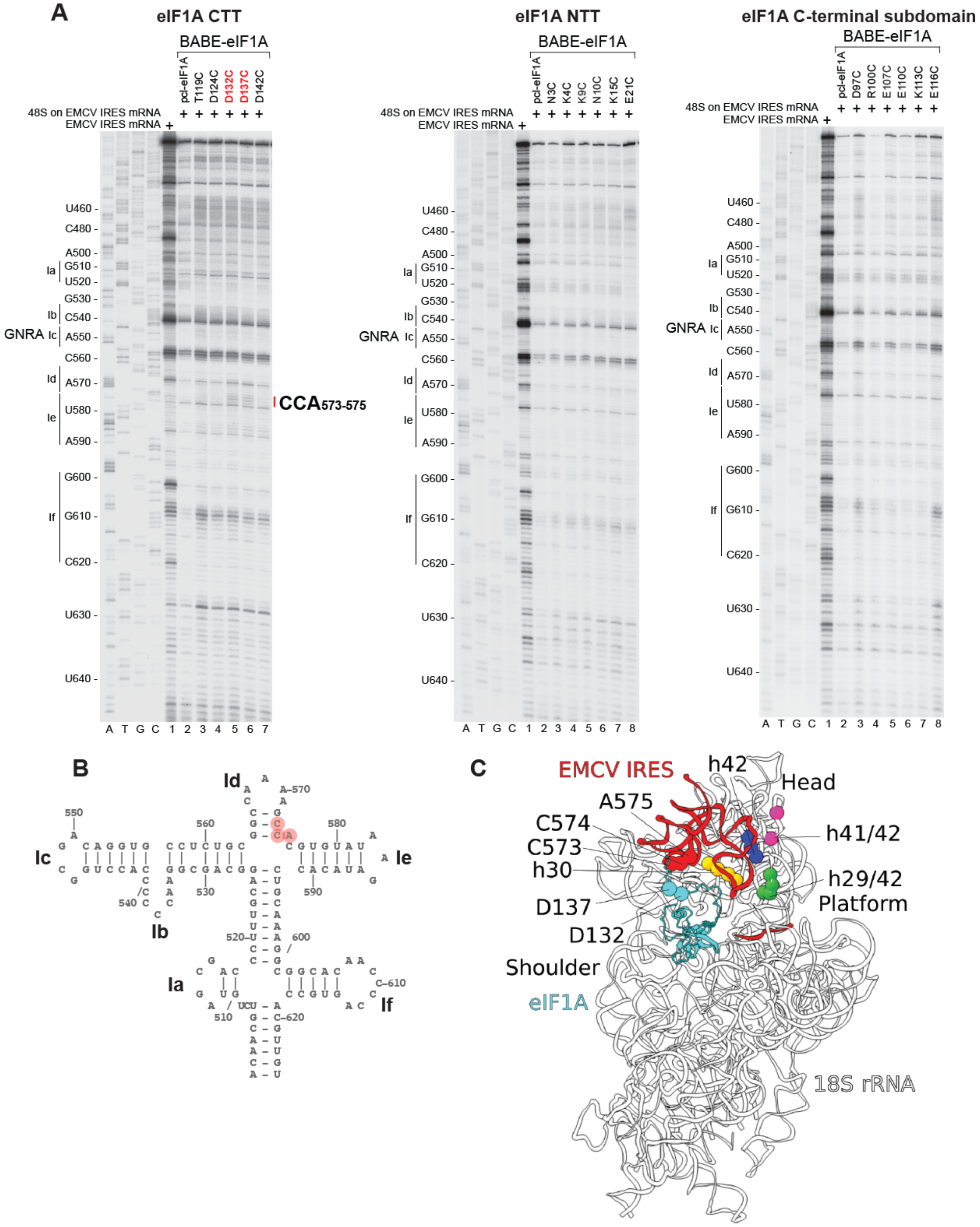


**Figure S5. Directed hydroxyl radical cleavage of the EMCV IRES in assembled 48S complexes from Fe(II) tethered to cysteines in eIF1A.**

(A) Analysis of directed hydroxyl radical cleavage of the EMCV IRES in 48S complexes from cysteines in the C-terminal tail (CTT) (left panel), N-terminal tail (NTT) (middle panel) and C-terminal subdomain (right panel) of eIF1A. Sites of cleavage were mapped by primer extension inhibition. Positions of cleaved nucleotides are shown on the right. Lanes G, A, T, C depict EMCV sequence generated from the same primer. (B) Sites of directed hydroxyl radical cleavage in the EMCV IRES from D132 and D137 in eIF1A mapped onto the secondary structure of the apex of domain I. (C) Position of directed hydroxyl radical cleavage in the EMCV IRES mapped onto the cryo-EM structure of the 48S complex and positions of nucleotides in 18S rRNA that are cleaved from the same residues of eIF1A (D132 and D137) in 43S preinitiation complexes[55].

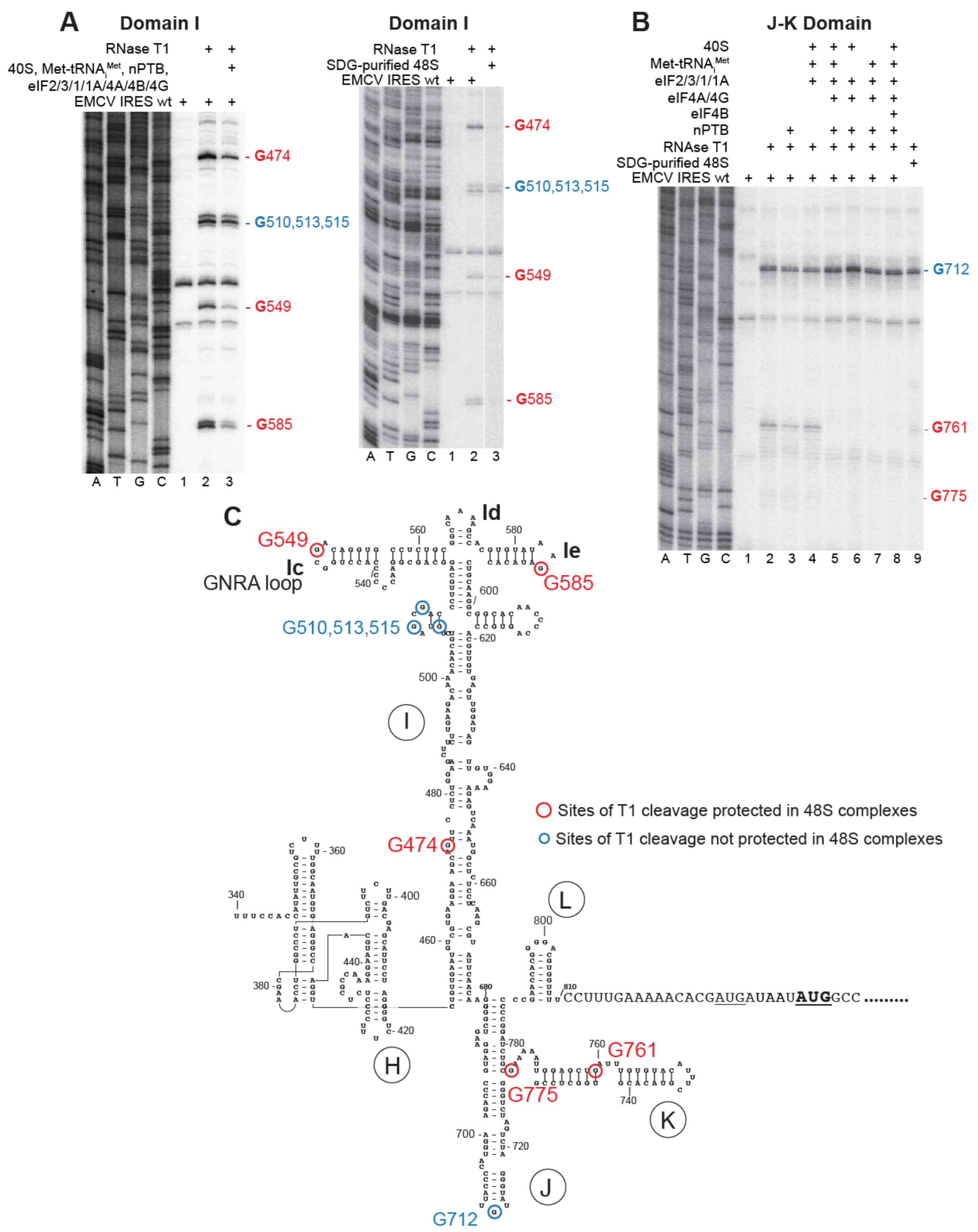


**Figure S6. Protection of the EMCV IRES from RNAse T1 cleavage in 48S initiation complexes.** (A) RNase T1 foot-printing of domain I in unpurified (left panel) and sucrose density gradient (SDG) purified (right panel) 48S complexes assembled on the EMCV IRES mRNA. (B) RNase T1 foot-printing of the JK domain in unpurified and SDG-purified 48S complexes assembled on the EMCV IRES mRNA. Sites of RNase T1 cleavage are marked on the right. Lanes C, T, A, and G depict EMCV sequence generated using the same primer. The division between lanes 2 and 3 (A, right panel) indicates that these two sets of lanes were derived from the same gel. (C) The EMCV IRES secondary structure model showing sites protected from the RNase T1 cleavage in 48S complexes.

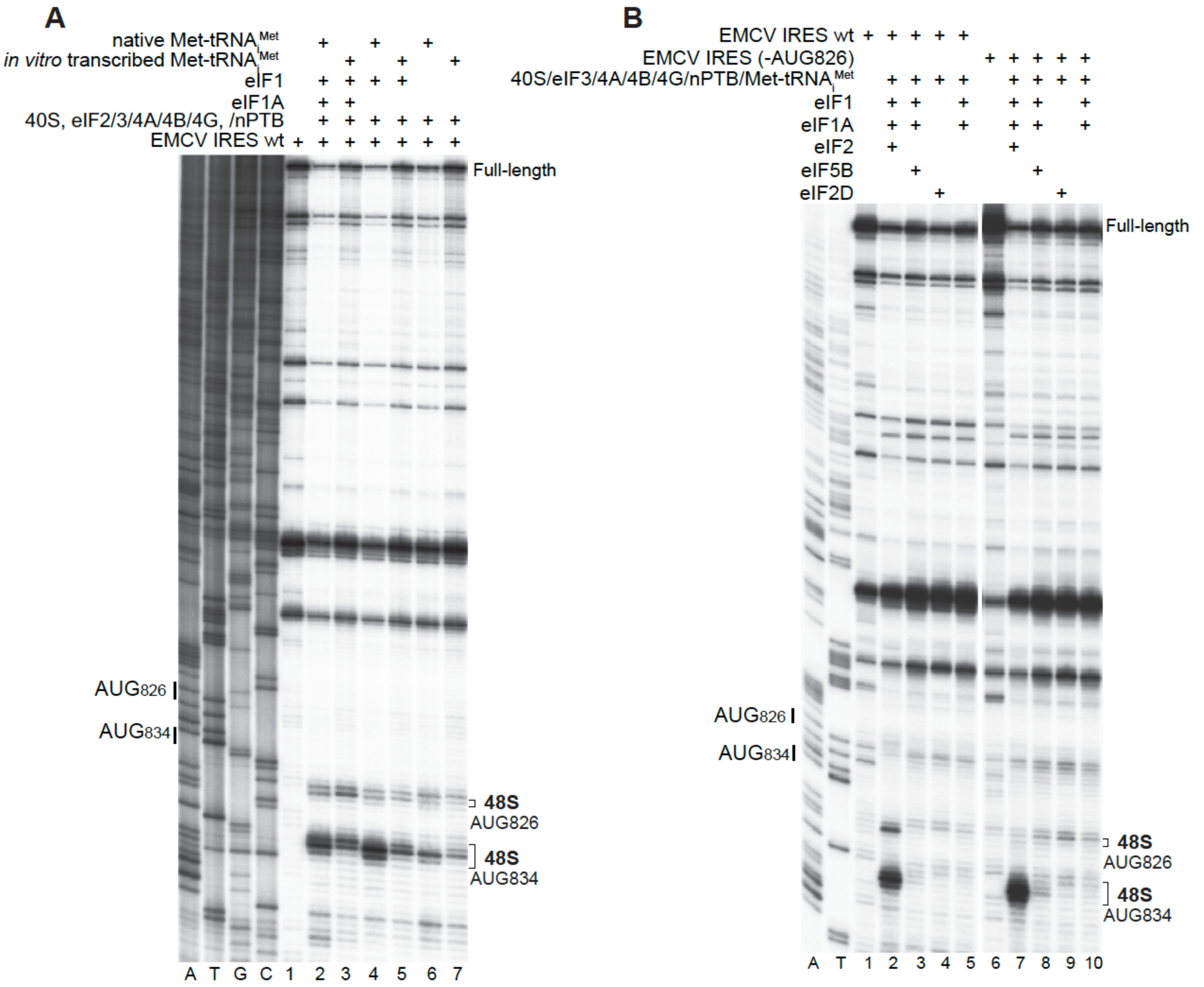


**Figure S7. Influence of the nature of initiator tRNA and substitution of eIF2 by eIF5B or eIF2D on 48S complex formation on the EMCV IRES.**

(A) Toe-printing analysis of 48S complex formation on wt EMCV IRES mRNA in the presence of either native or in vitro transcribed initiator tRNA, 40S subunits, nPTB and indicated initiation factors. (B) Toe-printing analysis of 48S complex formation on wt EMCV IRES mRNA and mRNA with mutated $AUG_{826}$ in the presence of 40S subunits, initiator tRNA, nPTB and indicated initiation factors. Lanes C, T, A, and G depict wt EMCV sequence generated using the same primer. The positions of initiation codons are indicated on the left and of assembled 48S complexes on the right. The division between lanes 5 and 6 (panel B) indicates that these two sets of lanes were derived from the same gel.

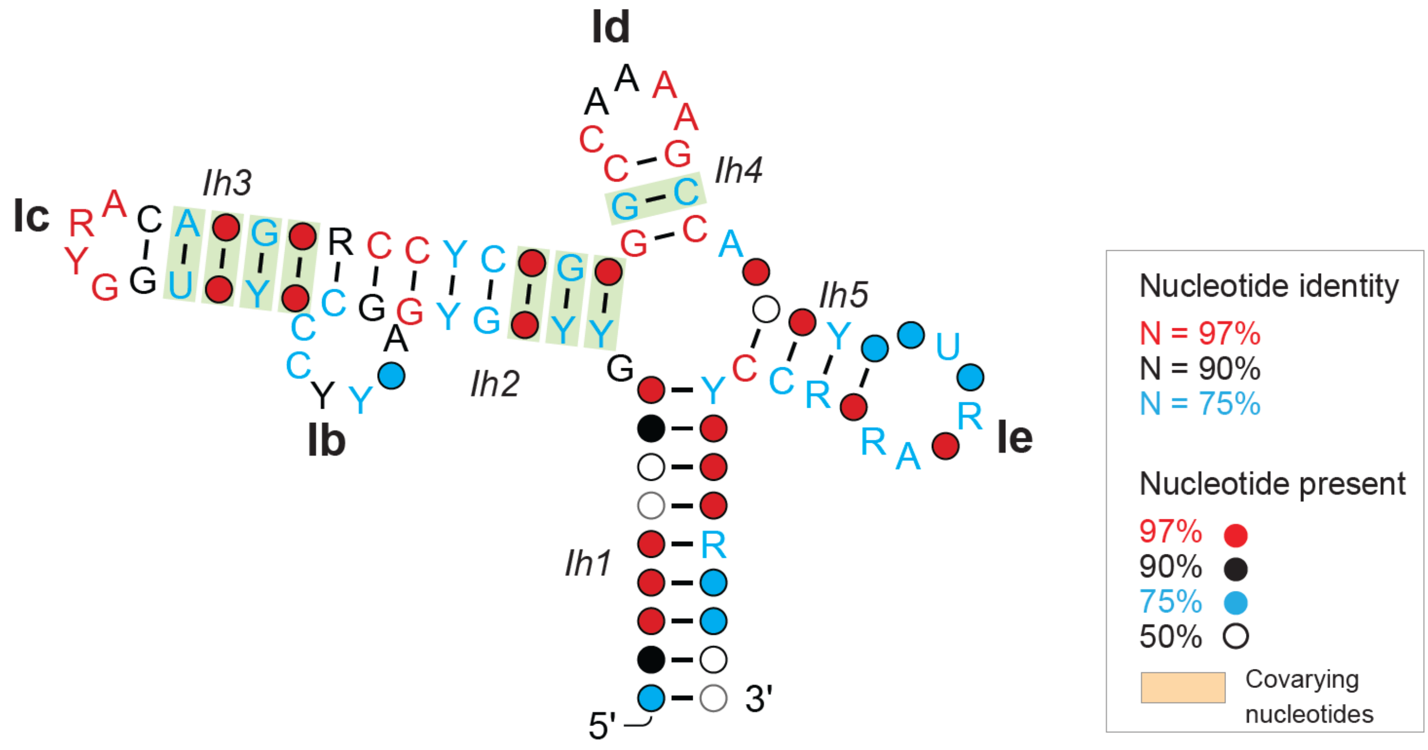


**Figure S8. Consensus model of the apical region of domain I of type 2 IRESs.**
The covariance model was derived using R-Scape and CaCoFold[87,88] using 89 curated picornavirus sequences (see Table S3), and summarized as a secondary structure with significantly covarying positions indicated with a green highlight. Y = U or C, R = A or G. The model is annotated to indicate structural elements.

**Table S1. Data collection statistics.**

| Parameter | Value |
|---|---|
| Microscope | FEI Titan Krios |
| Voltage | 300 kV |
| Detector | Gatan K3 Summit |
| Energy Filter | Gatan BioQuantum, 20 eV slit width |
| Magnification | 105,000× |
| Pixel size | 0.83 Å |
| Electron dose | ~60 $e^{-}$/Å$^2$ (total) |
| Exposure time | 2.5 s (fractionated into 50 frames) |
| Defocus range | –0.8 to –2.5 μm |
| Number of micrographs collected | 15,446 |
| Number of particles extracted | 835,875 |
| Number of particles after 2D classification | 688,588 |
| Symmetry imposed | C1 |
| Final map resolution (FSC 0.143) | 3.1 Å |
| Map sharpening B-factor | –85 Å$^2$ |
| Software used | cryoSPARC, RELION, Phenix, COOT |
| Model building & refinement | Phenix, Coot |
| Model resolution (FSC 0.5, masked) | 3.3 Å |

| | |
|---|---|
| MolProbity score | 1.75 |
| Clashscore | 6.8 |
| Ramachandran favored (%) | 96.2 |
| Ramachandran outliers (%) | 0.1 |
| Rotamer outliers (%) | 0.5 |
| CC (map-to-model) | 0.84 |

**Table S2. Search terms used to map visible nucleotides in the cryoEM structure of the 48S initiation complex assembled on the EMCV IRES to its nucleotide sequence.**

| Direction | Search terms | Hits | Possible |
|---|---|---|---|
| 5 to 3 | _RNRR | 39 | 2 |
| 5 to 3 | _RYRR | 23 | 1 |
| 5 to 3 | _RYRRNNNNNNNNYY | 9 | 1 |
| 5 to 3 | _RYRRYNNNNNNNYY | 6 | 1 |
| 5 to 3 | YRYRRNNNNNNNNYY | 5 | 1 |
| 5 to 3 | YRYRRYNNNNNNNYY | 3 | 1 |
| 5 to 3 | YRYRRNNNNRNNYY | 1 | 1 |
| 3 to 5 | RRNR_ | 42 | 2 |
| 3 to 5 | RRYR_ | 18 | 1 |
| 3 to 5 | YYNNNNNNNNRRYR_ | 7 | 1 |
| 3 to 5 | YYNNNNNNNNRRYRY | 3 | 1 |
| 3 to 5 | YYNNNNNNNYRRYRY | 0 | 0 |
| 3 to 5 | YYNNRNNNNRRYRY | 0 | 0 |

### Table S3. The apical region of domain I of picornavirus type 2 IRESs

| Genus | Virus | Acc. No. | Nts. |
|---|---|---|---|
| Ailurivirus | Ailurivirus D isolate QlGpf124Aim01-12 | MZ357174.1 | 886-968 |
| Aphthovirus | Foot-and-mouth disease virus A isolate abrazil iso67 | AY593788.1 | 779-857 |
| | Foot-and-mouth disease virus A isolate Hafizabad/QOL-UVAS-Pak/2005 | KY446902.1 | 769-847 |
| | Foot-and-mouth disease virus A isolate Egypt/El Minia/36/2016 | ON168597.1 | 58-135 |
| | Foot-and-mouth disease virus A isolate a3mecklenburg iso81 | AY593776.1 | 737-815 |
| | Foot-and-mouth disease virus SAT 2 isolate BOT-BUFF/107/72 | MH053331.1 | 735-811 |
| | Foot-and-mouth disease virus A isolate MCH/1557/2001 | OR338615.1 | 747-824 |
| | Foot-and-mouth disease virus Asia1/PAK/ICT/220-4/2012_pro | OM471639.1 | 774-852 |
| | Foot-and-mouth disease virus (FMDV) strain C, isolate c-s8c1 | AJ133357.1 | 726-804 |
| | Foot-and-mouth disease virus A isolate A/ALG/3/2017 | MG923580.1 | 244-321 |
| | Foot-and-mouth disease virus O isolate O1/Manisa/TUR/69 | KY825719.1 | 199-277 |
| Bopivirus | Bopivirus sp. strain goat/AGK16/2020-HUN | MW298058.1 | 236-315 |
| | Bopivirus sp. strain deer/VIC82-2020/AUS | MZ436972.1 | 301-378 |
| Cardiovirus | Boone cardiovirus 1 isolate BCV-1 | NC_038305.1 | 1114-1194 |
| | Cardiovirus A isolate 22084x5-1820 | MZ544194.1 | 252-334 |
| | Cardiovirus B strain rat08/rCaB/HUN | MN116646.1 | 753-834 |
| | Cardiovirus B isolate YNMIX-YNSC213 | PQ678021.1 | 650-732 |
| | Cardiovirus F1 isolate RtMruf-PicoV/JL2014-1 | NC_075977.1 | 499-580 |
| | Vole cardiovirus isolate 14657 | ON584156.1 | 576-658 |
| | Encephalomyocarditis virus | NC_001479.1 | 518-600 |
| | Encephalomyocarditis (EMC) virus EMC-D variant | M22458.1 | 513-595 |
| | Mengo virus isolate Rz-pMwt | DQ294633.1 | 437-519 |
| | Mengo virus strain AnrB-3741 | KU955338.1 | 230-312 |
| | Rat theilovirus 1 strain RTV-1 | EU542581.1 | 742-823 |
| | Cardiovirus B strain Ruian-Rn93-3 | MF352411.1 | 179-260 |
| | Cardiovirus B3 isolate 1 | MF172923.1 | 757-837 |
| | Genet fecal theilovirus isolate S15 | KF823815.1 | 1114-1194 |
| | Theiler's murine encephalomyelitis virus GDVII | X56019.1 | 751-833 |
| | Marmot cardiovirus strain HHMCDV | MZ382838.1 | 740-821 |
| | Cardiovirus C3 strain Wencheng-Rn416 | MF352424.1 | 1000-1081 |
| | Saffold virus strain Can112051-06 | JF813004.1 | 738-819 |
| | Saffold virus strain Penang | HQ162476.1 | 755-836 |
| | Saffold virus: isolate: Pak-2491 | AB747249.1 | 734-815 |
| | Saffold virus strain Nijmegen2008 | FN999911.1 | 735-816 |
| Cosavirus | Cosavirus A isolate MR96-15-1/GER/2015 | MT094345.1 | 245-327 |
| | Cosavirus A strain AM326/BRA-AM/2017 | MT023104.1 | 850-932 |
| | Cosavirus B isolate A21_AFP15_NGR_2020-B | PP386518.1 | 260-342 |
| | Cosavirus E isolate A28_AFP12_NGR_2020-E | PP386515.1 | 640-720 |
| | Human cosavirus isolate SEWAGE/NL/1999-046-1 | KJ437094.1 | 223-305 |
| | Human cosavirus A19 strain PK6187 | JN867759.1 | 45-125 |
| Erbovirus | Equine rhinitis A virus strain PERV-1 | DQ272578.1 | 622-699 |
| | Equine rhinitis A virus strain Plowright | DQ272127.1 | 278-355 |
| | Erbovirus A strain 303 | KX260138 | 547-623 |
| | Erbovirus A strain 396 | KX260139.1 | 550-626 |
| | Erbovirus A strain 421 | KX260140.1 | 547-627 |
| | Equine rhinitis B virus 1 | NC_003983 | 543-621 |
| | Equine rhinitis B virus 2 strain 1228 | KX260141.1 | 555-631 |
| | Equine rhinovirus 3 strain P313/75 | AF361253 | 555-631 |
| | Crocidura shantungensis picorna-like virus 5 isolate picorna_8 | PP272660.1 | 343-423 |
| | Picornavirales sp. isolate 193-k141_280731 | MZ678985.1 | 440-519 |
| | Wufeng shrew picornavirus 3 isolate WF_Cr.attenuata_picorna_2 | OQ716066.1 | 7--781 |
| Hunnivirus | Hunnivirus A9 isolate RtRrs-PicoV/YN2014 | KY432925.1 | 344-422 |
| | Hunnivirus A isolate JM_Ap.agrarius_picorna_1 | OQ715979.1 | 369-449 |
| | Hunnivirus A7 isolate 05VZ-75-RAT099 | KT944214.1 | 271-351 |
| | Hunnivirus A isolate LQ_Ra.tanezumi_picorna_1 | OQ715984.1 | 366-446 |
| | Pangolin hunnivirus isolate ZJ-MO7 | OM451179.1 | 404-483 |
| | Bovine hunnivirus strain BoHuV-WZ-202 | OQ790152.1 | 252-330 |
| | Ovine hungarovirus OHUV1/2009/HUN | HM153767.3 | 389-469 |
| | Porcupine hunnivirus isolate FJ-F1 | OM451178.1 | 345-425 |
| Mischivirus | Mischivirus sp. isolate MSWZC17/6 | OR867092.1 | 1085-1164 |
| | Cabezo Gordo bat-associated mischivirus | PP654855.1 | 1106-1185 |

| | | | |
|---|---|---|---|
| | Miniopterus schreibersii picornavirus 1 | NC_034381.1 | 1093-1172 |
| | African bat icavirus A isolate PREDICT-06105 | KP100644.1 | 835-912 |
| | Pteropus rufus mischivirus isolate AMB150 | OQ818316.1 | 829-910 |
| | Miniopterus bat picornavirus isolate 2A/Kenya/BAT0738/2015 | PP711935.1 | 1107-1186 |
| | Canine picornavirus isolate A128thr polyprotein (QKD15_gp1) | NC_075428.1 | 573-651 |
| Mosavirus | Mosavirus sp. isolate YSS01 | MW826550.1 | 336-409 |
| | Mosavirus A2 strain SZAL6-MoV/2011/HUN | NC_023987.1 | 352-425 |
| Parechovirus | Sebokele virus 1 | NC_021482.1 | 418-495 |
| | Ljungan virus strain 87-012G | EF202833.1 | 422-501 |
| | Ljunganvirus 5 | LC133331.1 | 409-487 |
| | Bovine parechovirus cow/2018/4 | BR001751.1 | 376-458 |
| | Bovine parechovirus Bo_Par/Den1/2021/JPN | LC650808.1 | 376-458 |
| | Parechovirus C isolate 22057x67-9 | MZ544294.1 | 117-197 |
| | Parechovirus E1 isolate falcon/HA18-080/2014/HUN | KY645497.1 | 444-522 |
| Rabovirus | Rabovirus A strain Wencheng-Rt38-1 | MF352417.1 | 408-486 |
| Rosavirus | Rosavirus M-7 | NC_038880.1 | 163-247 |
| | Rosavirus B isolate YY4 | PQ045668.1 | 51-134 |
| | Rosavirus B isolate RVB/YY86 | OM492423.1 | 151-232 |
| | Rosavirus B isolate RVB/YY106 | OM492427.1 | 201-282 |
| | Rosavirus B strain rat08/rRoB/HUN | MN116648.1 | 235-316 |
| | Rosavirus B isolate RVB/SZ59 | OM492432.1 | 139-221 |
| | Rosavirus C strain RASM14A | KX783433.1 | 373-455 |
| | Rosavirus C strain NFSM6F | KX783428.1 | 207-291 |
| Sapelovirus | Coypu sapelovirus 1 isolate GX-F1 | OM451188.1 | 420-495 |
| | Coypu sapelovirus 2 isolate HuN-A2 | OM451191.1 | 450-526 |
| Unclassified | Picornavirales sp. isolate 193-k141_280731 | MZ678985.1 | 440-519 |
| | Riboviria sp. isolate flycatcher172_contig_428 | OQ424099.1 | 6689-6608 |
| | Picornaviridae sp. isolate xizangnaqu19-7207 | OR367578.1 | 443-521 |
| | Picornaviridae sp. isolate YSN02 | MW826507.1 | 263-335 |

Names, accession numbers and inclusive nucleotide numbers of sequences from the named picornaviruses, belonging to genera as indicated, that belong to the apical region of domain I. Sequences were analyzed using R-scape; Cacofold was then used to generate a consensus secondary structure (Figure S8).
